# Generalized correlation-based dynamical network analysis: a new high-performance approach for identifying allosteric communications in molecular dynamics trajectories

**DOI:** 10.1101/2020.06.18.160572

**Authors:** Marcelo C. R. Melo, Rafael C. Bernardi, Cesar de la Fuente-Nunez, Zaida Luthey-Schulten

## Abstract

Molecular interactions are essential for regulation of cellular processes, from the formation of multiprotein complexes, to the allosteric activation of enzymes. Identifying the essential residues and molecular features that regulate such interactions is paramount for understanding the biochemical process in question, allowing for suppression of a reaction through drug interventions, or optimization of a chemical process using bioengineered molecules. In order to identify important residues and information pathways within molecular complexes, the Dynamical Network Analysis method was developed and has since been broadly applied in the literature. However, in the dawn of exascale computing, this method is generally limited to relatively small biomolecular systems. In this work we provide an evolution of the method, application and interface. All data processing and analysis is conducted through Jupyter notebooks, providing automatic detection of important solvent and ion residues, an optimized and parallel generalized correlation implementation that is linear with respect to the number of nodes in the system, and subsequent community clustering, calculation of betweenness of contacts, and determination optimal paths. Using the popular visualization program VMD, high-quality renderings of the networks over the biomolecular structures can be produced. Our new implementation was employed to investigate three different systems, with up to 2.5 M atoms, namely the OMP-decarboxylase, the Leucyl-tRNA synthetase complexed with its cognate tRNA and adenylate, and the respiratory complex I in a membrane environment. Our enhanced and updated protocol provides the community with an intuitive and interactive interface, which can be easily applied to large macromolecular complexes.

## 1 Introduction

In the last few decades molecular dynamics (MD) simulations have become an indispensable tool for mechanistic analysis in structural biology. From its first applications, revealing the fluid-like interior of protein that result from the diffusional character of local atomic motion [1], to more recent applications simulating entire organelles [2], the information content generated by MD studies has grown rapidly. With the increase of system sizes [3] and the frequent use of enhanced sampling techniques [4,5], came the need for new and enhanced analysis tools, capable of extracting information from massive amounts of data and generating new insight. The most diverse approaches have been applied to identify system features that are relevant to its biological functions, including clustering algorithms [6,7], dimensionality reduction techniques [8], and a variety of strategies from the so-called “big-data” and “artificial intelligence” fields [9–11]. Developed just over a decade ago [12, 13], a particularly interesting technique that has recently become popular is the analysis of dynamical networks [14–16]. This technique has been employed to study how groups of atoms interconnect in “communities” [17], and also the allosteric signaling in tRNA:protein complexes [12,18], glutamine amidotransferase [19], and many other systems [16,20]. More recently, these methods have also been applied to identify how force propagates through mechanoactive biomolecules [21–26], a fundamental question in mechanobiology. Network analysis has been even use to guide atomic force microscopy (AFM) based single-molecule force spectroscopy (SMFS) experiments [27].

The analysis of networks and their properties has a long history, with applications in diverse fields such as engineering [28, 29], and social networks [30], and their approach to modelling molecular systems is particularly fruitful, leading to a rich field of research [8, 12, 19, 31]. Using MD simulations to extract dynamical features from biomolecules, from simple proteins to complexes, one can convert the atomic representation of the system into a “nodes-and-edges” representation that can then be analyzed much like any other graph [32, 33]. A key source of information is the partitioning of the network in subgroups (or communities) using algorithms such as Girvan—Newman’s [34], providing information on cooperative motion within a protein’s subdomains, or on residues that mediate communication between communities. Both are computationally challenging tasks, and can become very expensive as the size of the network grows.

Perhaps the greatest interest in using dynamical networks lies on how networks can be used to investigate allosteric signaling in biomolecular complexes [35, 36]. In most complexes a response to a specific stimuli is regulated in a coordinated manner. Such allosteric mechanisms have been found to play key roles in the functions of many proteins [15]. From a thermodynamics perspective, allosteric mechanisms result from changes in enthalpic and/or entropic interactions across a biomolecular complex [37, 38]. Since allosteric regulation of a protein’s functions does not necessarily depend on large conformation shifts [39], multiple advances in network analysis techniques applied to MD simulations have stayed away from locating high mobility elements, and have instead focused on sub-optimal path calculations [40], to identify redundant communication networks within molecular complexes, and on identifying residues central to communication pathways [19, 41]. These application evolved from depending on contacts between residues close in space [41], to utilizing correlations between the motions of neighboring residues [42].

Once a network of connected nodes has been created to represent a molecular system, multiple computational methods have been applied to calculate the path(s) that bridge allosteric site to target site, such as Dijkstra’s or Floyd Warshall’s algorithm [43, 44]. This computational approach to the problem has been target of research for over a decade [42, 45, 46], and lead to the creation of tools that identify information pathways that take signal to target [12, 32, 40].

Although difficult to obtain experimentally [14, 16], the knowledge of the atomic motions and their collective behavior in proteins is essential to the understanding of their biological function [47]. In MD simulations, correlation analysis techniques can be easily employed to investigate this behavior. Pearson correlation coefficients have been widely used in the analysis of MD simulation data. Despite being relatively cheap to calculate, Pearson correlation does not account for non-linear contributions to correlations, and fails to asses correlations in perpendicular motion of atoms [48]. To avoid said pitfalls, generalized correlation coefficients, using the well-known Shannon mutual information [49,50], have been employed in multiple areas of research [51], including MD simulations [46].

Here, we focus on extending the applicability of the network analysis methodology through a new interface and implementation, analyzing the reproducibility of results using replicas of targeted systems, and ultimately improving our ability to interpret results from the vast amounts of raw data gathered from MD simulations. The package described in this work allows users to analyze MD results in popular trajectory formats, such as DCD, TRR or CRD, with no pre-processing. The methodology presented here is entirely contained in Jupyter notebooks and Python modules, making it easy and practical to apply the techniques. Additionally, our package prepares input scripts for the popular VMD [52] software, allowing for practical publication-level GPU-accelerated ray-tracing rendering [53] of biomolecular images. A similar approach connecting Python packages calling VMD for imaging has been previously described [54].

To demonstrate the applicability of our software, we have investigated three different biological systems (see Fig. 1), selected to display a wide range of sizes (number of atoms) and biological contexts. The first system is a small enzyme, namely the Orotidine 5’-phosphate decarboxylase (OMP-decarboxylase). The second is a Leucyl-tRNA synthetase (LeuRS), a tRNA-bound protein responsible for the setting of the genetic code. The last system investigated here is the respiratory complex I, a multi-subunit transmembrane protein complex that is part of the respiratory chain of organisms ranging from bacteria to humans.

**Figure 1.**
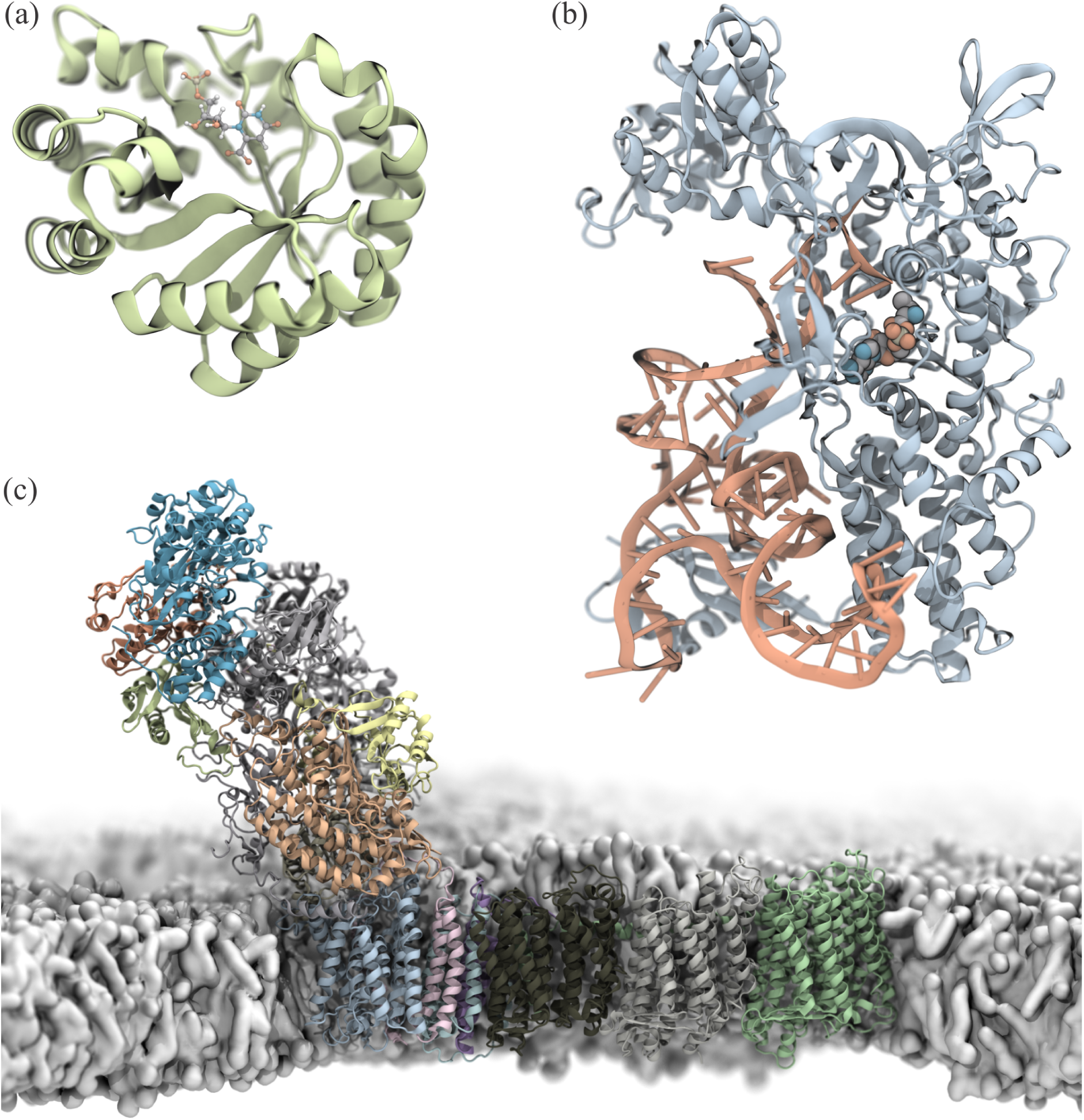
Biomolecular systems used to test the new Dynamic Network Analysis implementation. (a) Structure of a Orotidine 5’-phosphate decarboxylase monomer. (b) Strucutre of the Leucyl-tRNA synthetase. (c) Structure of the respiratory complex I embedded in a lipid membrane. All protein and nucleic acid structures are show in new cartoon representations, while ligands are shown in ball-and-stick representations. Images were rendered using VMD.

### 1.1 OMP-decarboxylase

The OMP-decarboxylase, also known as orotidylate decarboxylase, is a widely studied enzyme involved in pyrimidine biosynthesis. This enzyme catalyzes the decarboxylation of orotidine monophosphate (OMP), producing uridine monophosphate (UMP) [55]. The OMP-decarboxylase is probably the most efficient enzyme ever studied, reducing the energy barrier of the decarboxylation of OMP by several orders of magnitude [56]. The half-life of OMP in neutral aqueous solution is about 78 million years, but when catalyzed by OMP-decarboxylase its half-life is reduced to only 18 ms. Regarding its action mechanism, the enzyme is a member of the (*β*/*α*)8-barrel superfamily (see Fig. 1a), and its reaction mechanism has been thoroughly examined [57]. This system was chosen a small test system to create a tutorial (see Supplementary Material) for the new implementation of Dynamical Network Analysis presented here.

### 1.2 LeuRS complex

In order to translate genetic information, cells employ aminoacyl-tRNA synthetases (aaRSs) to charge tRNA with its cognate amino acid. This process is divided in two steps: an activation step, where the amino acid (or a precursor) reacts with an ATP molecule in the active site of the aaRS to form an aminoacyl-adenylate (aa-AMP); and a “charging” step, where the aaRS transfers the amino acid from the newly formed aminoacyl-AMP ligand to the adenine 76 base of its cognate tRNA. This is the essential process that assures the translation of genetic information into proteins. For each of the 20 naturally occurring amino acids, there is typically one aaRS tasked with charging it to its cognate tRNA [58, 59].

The *E. coli* Leucyl-tRNA synthetase (_*ec*_*LeuRS*, see Fig. 1b) is a class Ia synthetase, and its structure displays distinct sub-domains that have been extensively studied [60]. While many tRNA synthetases probe the tRNA anti-codon region to select the correct pair for amino acid binding, _*ec*_*LeuRS* lacks direct anticodon binding. It has been shown that _*ec*_*LeuRS* identifies the correct tRNAs using other identity elements. Indeed, several have been examined in the literature [61–64], highlighting a series of elements shared by the six different isoacceptors charged by LeuRS. Adenine 73 was observed to be an essential discriminator base for leucylation in multiple organisms, even leading to mischarging when a non cognate tRNA with a mutated A73 base was presented to LeuRS. Most strikingly, it was shown that a minimal RNA that had only its D-arm and T-arm intact (and its anticodon and variable loops deleted), could still be efficiently charged with leucine, indicating that those regions concentrated identifying information. Supporting this notion, bases G18 and G19 in the D-loop were found to be required for both recognition and aminoacylation. When G18:U55 and G19:C56 were experimentally mutated to maintain structural stability while exhibiting a different base pair, the aminoacylation activity was lost. Base A14, in the same D-loop, lead to a 100-fold reduction in aminoacylation when mutated, and its neighbour A15 also caused a drop in activity when mutated. For the *tRNA*^*leu*^(*UAA*) isoacceptor, U16 was shown to be important for aminoacylation [60], and only a pyrimidine mutation (C16) kept “reasonable” to native-like activity [62], however for the *tRNA*^*leu*^(*CAG*) form, the base was not essential for aminoacylation.

The thorough investigation of this tRNA:protein complex makes it a perfect target for detailed studies using Dynamic Network Analysis, and it will be the main focus of attention in this study.

### 1.3 Respiratory Complex I

The Respiratory Complex I is a large protein complex involved in the oxidative phosphorylation process, in which ATP is formed. Oxidative phosphorylation is the culmination of a series of energy transformations called cellular respiration or simply respiration in their entirety [65]. Although conceptually simple, the unraveling of the mechanism of oxidative phosphorylation has been one of the most challenging problems of biochemistry [66]. The flow of electrons through protein complexes located in the mitochondrial inner membrane leads to the pumping of protons out of the mitochondrial matrix [67]. The respiratory complex I, also known as NADH:ubiquinone oxidoreductase, is the first large protein complex of the cellular respiratory chain [68]. It catalyzes the transfer of electrons form NADH to coenzyme Q10 and translocates protons across the inner mitochondrial membrane [67, 69]. The proton translocation is performed by four proton pumps that operate in parallel, in a mechanism not well understood [70, 71].

The respiratory complex I is the largest of the respiratory complexes. Its structure is in an “L” shape with a long membrane domain and a hydrophilic domain, typically called peripheral-arm, where redox centers are found. Multiple experimentally determined structures are available, with a large variation in size from 500 kDa to over 1 MDa, depending on the organism where they are found. There are 14 strictly conserved core subunits that are necessary and sufficient for function. Here we chose the *Thermus thermophilus* complex, which has 16 subunits [72], in a membrane environment (see Fig. 1c), in order to test the limits of the new implementation our method. Our technique can be, in the future, used to understand how the electron transfer process in the peripheral-arm activates the transport of protons across the transmembrane domain of the respiratory complex I, which is one of the main open questions of the respiratory mechanism.

## 2 Theory

Network analysis is a relatively new and emerging field of science that bridges traditional social network analysis and link analysis within network theory. In molecular dynamics analysis, network analysis is typically performed using correlation of motion to determine the existence and strength of a link between different atoms or molecules of a system.

### 2.1 Generalized correlation coefficients

The generalized correlation coefficient is derived from a mutual information estimate *I*, calculated using the positions of a pair of nodes. *I*’s estimation, in turn, is based on an information theoretical approximation of Shanon’s entropy, as described in [48]. Briefly, the method takes two nodes *i* and *j* (representing two atoms), and determines *I* based on the number of simulation frames in which the nodes’ position vary less than a dynamic cutoff value given by a parameter *k* (see Eq. (1), which was obtained from Eq. (9) of [48]).

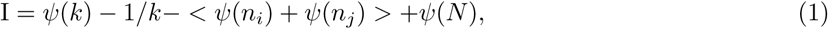

Here, *k* is set to 6, as proposed by Kraskov [48]. Additionally, *N* is the total number of simulation frames, *ψ*(*x*) = Γ(*x*)^*–*1^*d*Γ(*x*)*/dx* is the digamma function, and *n*_*i*_ and *n*_*j*_ are the number of frames in which the positions of nodes *i* and *j* are close to the one in a reference, and they are averaged by varying the reference frame over all simulation. The mutual information estimate is then transformed in a generalized correlation coefficient, by applying Eq. (2), where *d* = 3 for the (*x, y, z*) dimensions that describe each node.

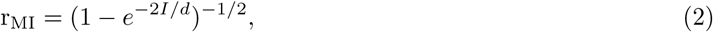

The calculation of mutual information estimate between a pair of nodes can be described more specifically as follows. Given a pair of nodes *i* and *j*, and a reference simulations frame *f*_0_, the position of each node is compared with their positions in all other simulation frames. For each frame, the highest of the variations in *x, y* and *z* dimensions is selected to represent the node’s “distance” from its position in frame *f*_0_ (note that the maximum variation among all dimensions is used, *not* the Cartesian distance). For each frame in the simulation, the highest of the distances for *i* or *j* is then selected, and used to sort all frames. Taking the *k* nearest neighbours in “simulation frame” space, meaning the *k* frames in which the positions of nodes *i* and *j* vary the least compared to frame *f*_0_, we can determine the maximum variation among the *x, y*, and *z* dimensions for each node, *i* and *j*, individually, giving *d*_*i*_ and *d*_*j*_. The two distances are used as cutoffs to select the frames where nodes *i* and *j* are closer than *d*_*i*_ and *d*_*j*_, with respect to their respective positions in frame *f*_0_. *n*_*i*_ and *n*_*j*_ are the number of simulation frames that meet these criteria. The same calculation is performed varying *f*_0_ from the first to the last frame, giving the values to the mean calculated in Eq 1. The generalized correlation coefficient is the achieved by applying the mutual information estimate in Eq. (2).

In this work, the calculation of a generalized correlation coefficient for a pair of atoms (or “nodes”), defined by their 3D positions over a series of MD simulation frames, was implemented in Python using elements of NumPy and Numba [73].

### 2.2 Dynamical network analysis

In dynamic network analysis, each residue of the biomolecular system being studied is represented by “nodes”. By default, amino acid residues are represented by a single node located in their alpha-carbons, and nucleotides by two nodes, one in the backbone phosphate, and one in the nitrogenous base. Water molecules have one node in their oxygen atom, and ions are trivially represented by one node. Adenylate residues are represented by three nodes, as they are the union of a nucleic base and an amino acid.

To determine which nodes are in contact, the shortest distance between heavy atoms (all atoms excluding hydrogen atoms) represented by two nodes is calculated. If the distance is shorter than 4.5°A in a simulation frame, the pair of nodes is said to be in contact in that frame. If a pair of nodes is in contact in more than 75% of a simulation, they are considered to be in contact for the purposes of network analysis. If one of the user-defined solvent molecules follows the contact criteria, these molecules are also included in the network.

The software package developed in this work uses MDAnalysis [74,75] to load and access atom position data from all MD simulations into the Jupyter notebook. Any popular trajectory format can be read, such as DCD, TRR or CRD, and no pre-processing is necessary. In order to obtain the best performance for contact detection and correlation calculations, parts of the code were optimized using Cython [76] and Numba [73]. Network statistics and determination of optimal paths were carried out using the Floyd-Warshall algorithm, provided by the NetworkX package [77]. For Floyd-Warshall calculations, the “distance” between nodes was defined as *d* = –*log*(*r*_*MI*_), consistent with previous applications of this method [12]. Community assignment was performed using the “Multilevel” algorithm [78], using the generalized correlation coefficients as edge weights.

All files necessary for automatically determining solvent and ion residues relevant for the analysis, calculating contact matrices, calculating generalized correlation coefficients in parallel for pairs of nodes in contact, and determining network properties like betweenness, clusters and shortest paths are available as supplementary material (in the form of Jupyter notebooks) and as a python package in the readily accessible Python Package Index (PyPI).

## 3 Simulation Details

Preparation of all simulations was done using VMD’s [52] QwikMD [79] interface. MD simulations were performed for all systems employing the NAMD [80] molecular dynamics package. The CHARMM36 [81] force field along with the TIP3 [82] water model was used to describe all systems. Simulations were carried out assuming periodic boundary conditions in the NpT ensemble with temperature maintained at 300 K using Langevin dynamics for pressure, kept at 1 bar, and temperature coupling. A distance cutoff of 12.0 °A was applied to short-range, nonbonded interactions, while long-range electrostatic interactions were treated using the particle-mesh Ewald (PME) method. The equations of motion were integrated using the r-RESPA multiple time step scheme to update the van der Waals interactions every two steps and electrostatic interactions every four steps [83]. The time step of integration was chosen to be 2 fs for all simulations performed. All analysis, other than network analysis, were performed using previously established protocols [84–86].

### 3.1 OMP-decarboxylase

The structure of the *Methanothermobacter thermautotrophicus* OMP-decarboxylase was obtained from the protein data bank (PDB), accession code 1X1Z [87]. Although the enzyme was crystallized as a dimer, the most biologically active form of the protein, for the purposes of the tutorial only one protein chain was simulated. It is important to notice that the monomer is still known to be enzymatically active. In order to obtain the co-crystal of the enzyme with its substrate, this substrate was modified to a 6-hydroxyuridine-5’-phosphate. Therefore, to be able to simulate the relevant system, we have modified the substrate employing VMD’s [52] Molefacture (see Fig. 1a). QwikMD was then employed to prepare a solvated system, with about 48,000 atoms, for simulation. Using NAMD, the system was minimized for 1,000 steps. Constrained MD simulations were performed for 1 ns, where the position of the protein backbone atoms, and ligand non-hydrogen atoms, were constrained by a Hook potential following standard NAMD protocols [83]. Then, 10 ns of unconstrained MD simulations were performed.

### 3.2 LeuRS complex

The structure for the ternary complex *LeuRS* : *tRNA*^*leu*^ : *Leu* –*AMP* was taken from the *Escherichia coli* crystal structure deposited under PDB ID 4AQ7 [60]. Chains A and B were used to create the ternary complex, and 8 unresolved nucleotides in two sets of 6 and 2 residues were modelled using RNA composer [88]. Magnesium ions were used to neutralize the system, and magnesium chloride was added in the simulation box to replicate experimental conditions. One nonstandard nucleotide was used in the tRNA. Uridine 16 was shown to be linked to catalytic activity [60], and was mutated to dihydrouridine in order to more closely model the system in its biological state.

The ternary complex *LeuRS* : *tRNA*^*leu*^ : *Leu* –*AMP* contained the cognate leucine adenylate (LeuAMP) in the active site (see Fig. 1b). Employing QwikMD the simulation system was prepared containing just over 260,000 atoms. Using NAMD, this complex was equilibrated (with position constraints for backbone atoms) for 50 ns in 50 independent replicas, and from their respective equilibrated structures, another 50 ns of simulation was used for analysis.

### 3.3 Respiratory Complex I

The complex I of respiratory chains plays a central role in cellular energy production. The respiratory chain is made by many transmembrane proteins that work in tandem. Here, the structure of *Thermus thermophilus* respiratory complex I was obtained from the structure deposited in the PDB under accession code 4HEA [72]. Gaps in subunit 6 were solved employing MODELLER 9.17 [89]. VMD [52] and its plugins (Molefacture and QwikMD) were employed to assemble the whole complex I system, including a POPC lipid membrane, as presented in Fig. 1c. The complete system comprises nearly 2,500,000 atoms, including 3,182 lipids. Using NAMD, the system was minimized for 5,000 steps. Then, 10 ns of constrained MD simulations were performed, where the position the protein backbone atoms were constrained by a Hook potential in a well described protocol [83]. An unbiased production run was performed with the GPU-accelerated NAMD for 100 ns. The trajectory from this last simulation was then employed in all the generalized network analysis presented here for the respiratory complex I.

## 4 Results and Discussions

Three systems with different levels of size and complexity were selected as a test bed for our new generalized network analysis software. The OMP-decarboxylase was used as our “tutorial system”, therefore little is presented in the main manuscript about this system. The small size of this enzyme and the fact that it is bound to its substrate, makes the OMP-decarboxylase a perfect example for those who want to learn how to use network analysis tools in their MD studies, particularly for those interested in drug development. The tRNA-bound enzyme LeuRS system was selected to be more carefully described in this manuscript. The third and last system was selected due to its complexity and for the fact that, like about a third of the human proteins, functions in a lipid membrane. The respiratory complex I will be used here mostly for its lipid-protein interactions, and to show how allosteric pathways evolve over time in an MD simulation.

### 4.1 OMP-decarboxylase

In the pharmaceutical field, there is a great interest in the interactions between enzymes and small molecules that work as substrates or inhibitors of these enzymes. Furthermore, very few enzymes can be considered as efficient as the OMP-decarboxylase [56, 90]. This widely studied enzyme was used by us as a sandbox for the development of our new network analysis tool. As such, we have produced a comprehensive tutorial, presented here as supplementary material. The tutorial allows the users of the generalized dynamical network analysis software to not only learn how to perform these analysis, but also how to include their own scripts to direct-target their research interests. The tutorial is divided in three parts, the first two using jupyter notebooks, and the last one using VMD. In the first part, all the “heavy-lifting” data processing is done, the generalized correlation calculation and all of the the most time-consuming steps are performed. The user will provide all the necessary information about the molecule, such as the ligands, and how to break the ligand in groups that are represented by multiple nodes. The main concepts regarding how network analysis works are introduced in this first part.

In the second part, the user is presented with another jupyter notebook where all the correlation maps calculated in the first part can be translated into molecular visualizations or plots. Here we provide an opportunity for users more comfortable with python programming to tailor the analysis to their specific scientific questions. In the third part, the user can load files produced by the Jupyter notebooks into VMD. An easy to load script will handle all the work, creating a simple graphical user interface (GUI) where the user can easily render publication-quality images. These renderings can represent many aspects of the biomolecular system. For instance, the tutorial will allow our users to easily produce a high-quality image of the full generalized network, as shown in Fig. 2a, or even how the communities and the betweenness of the biomolecular system are superimposed (see Fig. 2b). Communities and betweenness are dynamical network properties that will be better discussed along this manuscript. Fig. 2c shows how the OMP ligand interacts with the enzyme’s amino acid residues. Such analysis is particularly useful for those developing new drug molecules that may inhibit or act as substrate of an enzyme. For instance, here one of the most stable contacts was observe to be between OMP and Asp70, the amino acid that acts stabilizing OMP’s CO_2_, a key step in the enzymatic reaction of the OMP-decarboxylase. For more details on how to perform these analysis, see the tutorial presented as supplementary material. The tutorial was prepared to be easily adapted to other studies, particularly for those interested in drug-protein interactions.

**Figure 2.**
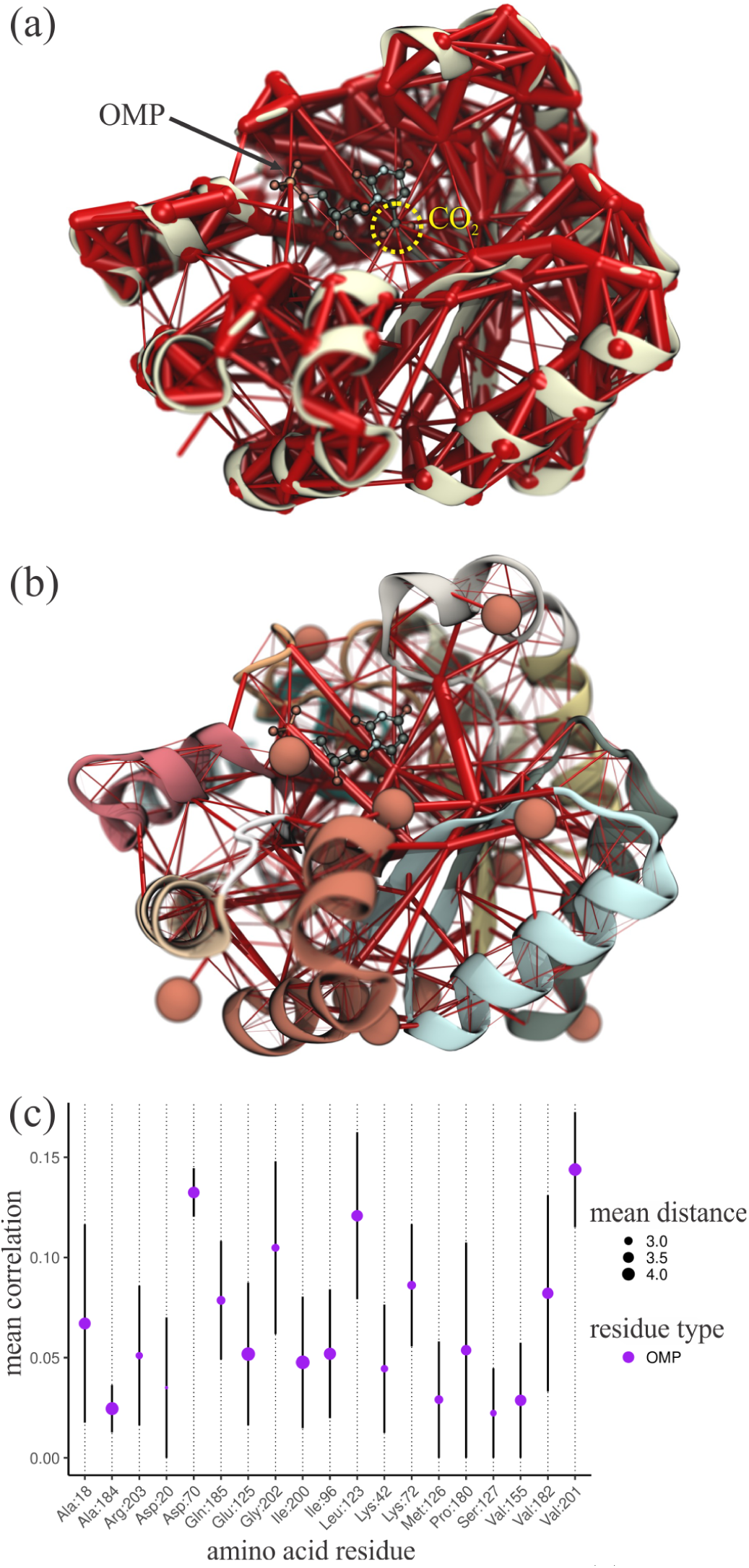
Analysis of the OMP-decarboxylase dynamic network. (a) Full network revealing the most correlated regions of the enzyme. The weight of the network edges (represented by thickness of red tubes) is given by its normalized generalized correlation coefficient. Correlation values are normalized from zero to one in order to produce the visual representations depicted in this and following figures. (b) Rendering showing communities and betweenness values of edges of the OMP-decarboxylase dynamic network. Communities are delineated by the different colors of the protein secondary structure, while betweenness values of network edges are indicated by the thickness red tubes. Both (a) and (b) images were rendered with VMD using our new Network Viewer 2.0 GUI. (c) Mean generalized correlation coefficients for contacts between OMP and amino acid residues in the OMP-decarboxylase active site. The *x* axis is labeled by amino acid residue, and the *y* axis indicates average generalized correlation coefficient (vertical black bars indicate standard error of the mean). Circle size indicates the average Cartesian distance between the closest heavy atoms in amino acids and OMP.

### 4.2 LeuRS complex

As the main application of our new generalized correlation-based network analysis tool, the LeuRS system bound to its cognate tRNA was used to not only test the capabilities of this new implementation, but also its performance. The protein was chosen for multiple reasons, including its evident biological relevance, being one of the enzymes that set the genetic code. Since the intent was to benchmark and showcase the generalized correlation-based network analysis, we also looked for a well described molecular system, and LeuRS has extensive experimental and computational literature previously dedicated to its study [60–63, 91]. Finally, the main application also needed to provide the opportunity to study intermolecular interactions, as opposed to just intra-molecular communication pathways. The LeuRS-tRNA-LeuAMP complex meets the demand with interfaces between different types of molecules: a protein, an RNA, and a small ligand.

#### 4.2.1 Performance

Using the LeuRS complex as a test system, 50 ns of MD trajectories in 50 independent replicas of the system were analyzed for node contacts. Approximately 1,150 nodes were studied per replica (the exact number varies since different amounts of water and ions were stably bound to the complex), and it was found that among all simulations, of the (*n* (*n* –1)*/*2) possible pairs of nodes in direct contact, only between 0.68% and 0.72% of the pairs were actually in contact. By constraining the expensive calculation of the generalized correlation coefficient to those pairs of nodes in direct contact, it was possible to avoid ∼99.3% of the computational effort.

More importantly, since determining the pairs of nodes in contact is an essential step in all dynamic network analysis studies, we can use the contact matrix to reduce the complexity of the calculation of generalized correlation coefficients (represented in the rendering in Fig. 3a) from ∼*O*(*N* ^2^) to ∼*O*(*N*) (see Fig. 3b,c). Considering an average number of contacts per node *c*, each node will participate in *c* *(*c* –1)*/*2 contact calculations (one calculation per unique pair of nodes), where *c* depends on the rules for determining a contact between nodes (see THEORY), not on the total number of nodes of the system.

**Figure 3.**
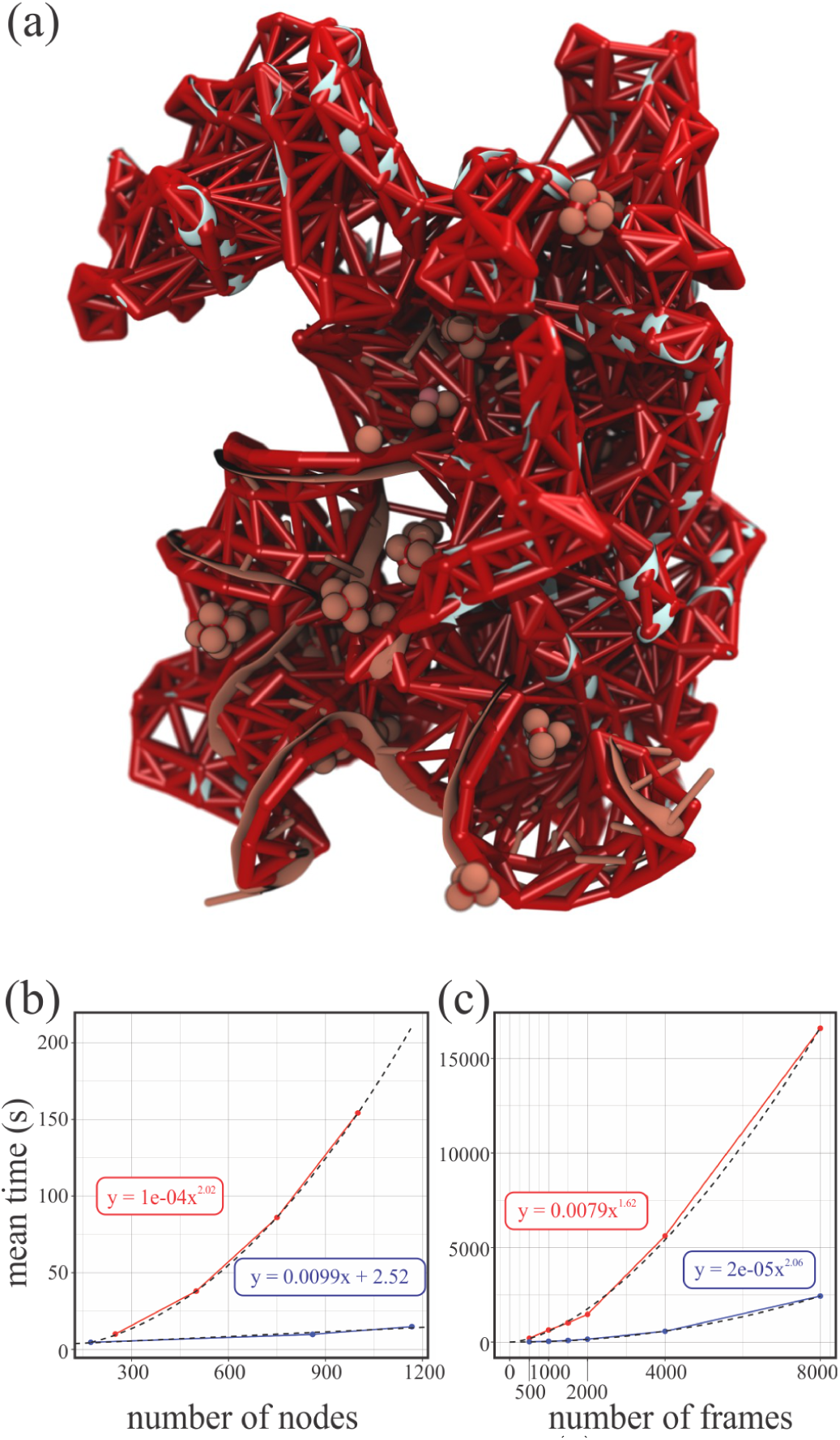
Generalized network analysis performance (a) Rendering of the LeuRS generalized correlation-based dynamic network. The full network reveals the most correlated regions of the enzyme-tRNA complex, where edge weights are given by their generalized correlation coefficients. (b) Scaling benchmarks for the calculation of generalized correlation coefficients. Average calculation time is shown as a function of to number of nodes in the system. In red, the cost of the calculation using the implementation described in [49]. The scaling is proportional to the square of the total number of nodes in the system. In blue, the linear scaling in the current implementation, which only calculates generalized correlation coefficients between nodes in contact. (c) Scaling for the calculation of the generalized correlation coefficients with respect to sampled simulation frames. All benchmark calculations were carried out using a dual Xeon E5-2650 v2 CPU with 2.6 GHz, and 32 GB of RAM.

To benchmark the current implementation (see Fig. 3b,c), subsets of the binary system *LeuRS* : *tRNA*^*leu*^ were analyzed, using either the tRNA, the protein, or the tRNA:protein complex, totalling 174, 860 and 1034 nodes respectively. The number of unique node pairs in contact was 444, 3,850 and 4,816, respectively. For the calculation using the method described in [49], since no contact determination is necessary, progressively larger number of nodes were used for the benchmark disregarding the spacial distribution of said nodes. Groups of 250, 500, 750 and 1,000 nodes were used, and all pairwise generalized correlation coefficients were calculated, totalling 31,125, 124,750, 280,875, and 499,500 unique node pairs, respectively.

Analyzing the 50 independent replicas of the ternary complex, it was found that nodes had approximately 8.25 direct contacts over the whole system. Table 1 lists the average number of contacts per node, considering the different types of residues they represent.

**Table 1.** Average number of contacts per node type. Contacts from nucleotide nodes are averaged between phosphate backbone and nitrogenous base nodes. Average contacts were calculated averaging the number of contacts of all nodes of the same type in one MD simulation. The average contacts for adenylate nodes were averaged across the 50 replicas since only one Leu-AMP molecule exists per replica.

| Type of Node | average contacts | Number of nodes |
| --- | --- | --- |
| All | 8.25 | 1167 |
| Amino acid | 9.0 | 860 |
| Nucleotide | 6.2 | 174 |
| Water | 5.7 | 111 |
| Ion | 6.1 | 19 |
| Leu-AMP | 10.3 | 3 |

With the accelerated generalized correlation calculation, the main time constraint to applying dynamic network analysis becomes calculating the contact matrix for the system in question. We see a slight drop in performance when comparing the current implementation (based in MDAnalysis and Cython optimized functions) with the original implementation [12] (see Fig.3), but simple mitigation strategies can be adopted to avoid a significant drop in performance for large systems.

Estimating mutual information using the Kraskov *et*.*al*. [48] method depends on re-ordering simulation frames, therefore sorting strategies will have a large impact on the efficiency of the method as larger simulations are used to calculate correlations. Fig. 3b shows the scaling of the current and the Lange [49] implementations when calculating generalized correlation coefficients for the LeuRS ternary complex. It is clear from the curve fits that the elaborate sorting strategy chosen in [49] scales better with respect to number of frames, but since it still relies on correlation calculations between *all* nodes, the overall cost is much higher than that of the current implementation. We note that the current implementation also allows for parallelization.

With the improvement in performance for calculation of correlation coefficients, the network analysis framework becomes constrained at an earlier stage: the determination of a contact matrix. As the contact matrix describes which nodes are connected to which other nodes in the system, it will invariably depend on a “all-to-all” distance calculations (an *O*(*N* ^2^) process), in order to determine which residues are closer than a cutoff distance from a reference residue. This calculation becomes very expensive very quickly, and a large amount of frames (many hundreds to thousands) from an MD simulation must be sampled as to create a good contact matrix.

One approach that could help mitigate this issue would be to perform the contact detection in multiple stages. In an initial stage, in order to estimate the nearest neighbours of a reference residue, a larger cutoff distance is used, and a smaller number of sampled frames is scanned. In a following stage, the distance cutoff is lowered, and more frames are used to detect contacts, but only distances between the reference residue and its neighbours from the previous stage are calculated. At each stage, the list of neighbours is trimmed, and number of distance calculations will be smaller. The sub-sampling of frames from the MD trajectory would need to be done carefully, as to avoid missing contacts.

Since the limitation created by the contact matrix creation is considerably smaller than that of the calculation of generalized correlation coefficients, the problem was not directly tackled in this work. However, the mitigation strategy proposed above was used in out tutorial and can be extended in case contact detection became a limiting factor in larger systems.

As for the cost of re-ordering simulation frames while estimating the mutual information between tow nodes, there are several fast sorting strategies available in widely used software packages. Here, the “quicksort” method implemented in NumPy was chosen for its ease of use and good performance. It is worth mentioning that,as simulations grow longer, it will be more informative to cut large trajectories into windows and analyze the system’s progression over different states, instead of “averaging” large conformational changes into one contact matrix and correlation network. Therefore, even for long MD simulations, the number of frames in each window would be relatively small, keeping this factor from hindering the calculation of correlations.

The trade-off between keeping “large enough” windows that will reliably capture system’s state, and “small enough” windows that will not average out important fluctuations, is not something we believe to be system *independent*, and requires careful case-by-case examination.

#### 4.2.2 Generalized network communities within the *LeuRS* : *tRNA*^*leu*^ complex

Another common way of analyzing dynamic networks is by looking at the communities formed by the network nodes. In molecular systems, such analysis is helpful in the identification of protein domains and how they interact with one another. In dynamic networks, a network is said to have community structure if the nodes of the network can be grouped into sets of nodes that are internally well-connected. The approach indicates that the network is subdivided into groups of nodes with dense connections internally and sparser connections between groups. Therefore, pairs of nodes are more likely to be connected if they are both members of the same community, and less likely to be connected if they are not.

Finding communities within an arbitrary network is frequently a very computationally demanding task. Despite the difficulties in finding these communities, several methods have been developed and employed with varying levels of success. Here, we have used the Louvain heuristics [78] to investigate how the nodes of the LeuRS complex are group into communities. The Fig. 4a reveals that the system is subdivided in a handful of communities, with the tRNA becoming mostly part of three different communities. It is interesting to note that there are few communities grouping tRNA and LeuRS nodes. Fig. 4b also shows the weaker edges connecting communities to one another.

**Figure 4.**
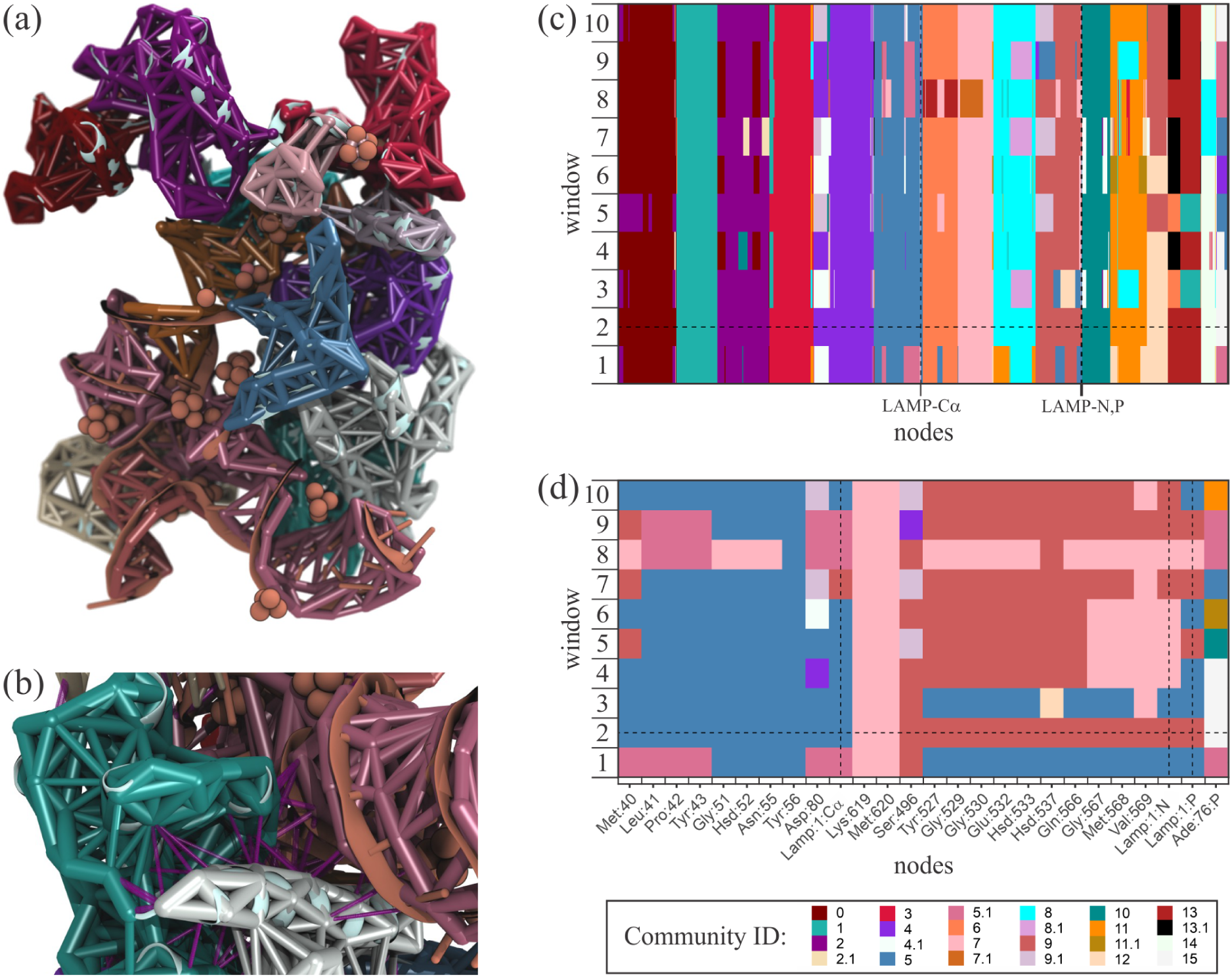
Rendering of the LeuRS generalized correlation-based communities. (a) Different communities are represented by different colors of the nodes and edges in the network. (b) In a different point of view, the inter-community links are shown as purple dashed lines. (c) Community statistics from different independent replicas (or “windows”) of the same system. Nodes were grouped by community as to highlight the high persistence of the assignment of most amino acid and nucleic acid residues to the same communities across independent replicas. (d) Same as (c) for residues in the catalytic site which make direct contacts with the adenylate.

The Louvain heuristics method was chosen here because it outperforms all other commonly used methods when accounting for speed and accuracy, as thoroughly investigated by Yang *et. al*. [92]. In particular, it consistently outperforms competing methods within the range of network sizes one tends to encounter in network analysis of macromolecular complexes (hundreds to thousands of nodes).

#### 4.2.3 Determination of identity elements in the LeuRS interface

The ternary complex LeuRS was studied in 50 independent MD simulations, and the results were analyzed with the updated Dynamic Network Analysis framework. Out of more than 250 interface contacts (direct contacts between protein and tRNA or adenylate), our method could identify identity elements in the protein:tRNA interface that guarantee the correct binding of the protein to its cognate tRNA (see Fig. 5).

**Figure 5.**
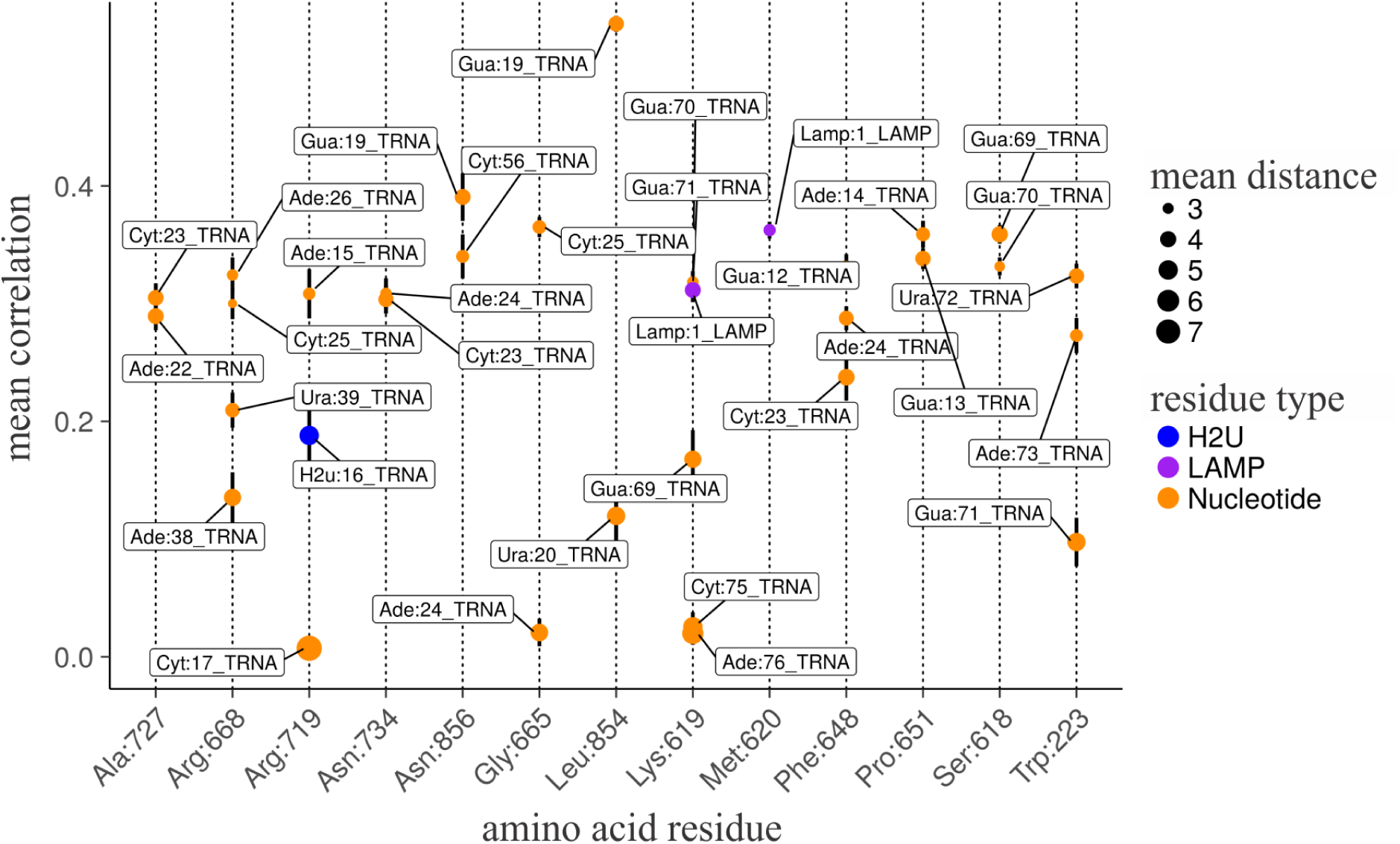
Mean generalized correlation coefficients for contacts along the protein:tRNA interface. The figure shows pairs of nodes along the protein:tRNA interface. The *x* axis is labeled by amino acid residue, and the *y* axis indicates average generalized correlation coefficient (vertical black bars indicate standard error of the mean). Labels indicate the second node in the interacting pair, circle size indicates the average Cartesian distance, and colors discriminate the type of residue (LAMP is the Leucyl-AMP adenylate, and H2U is the modified dihydrouridine 16). We show amino acid residues which have at least one connection with a mean correlation higher than 0.3.

Several experimentally verified identity elements show clear binding patterns in our simulations. Arg719 is part of the conserved K/DD/RR motif, and makes contacts with the modified base dihydrouridine 16, as well as neighboring bases A15 and C17. D16 was shown to be essential for binding and for catalytic activity [60]. Both G19-C56 and A14-U48 are known base pairs shown to be important for both recognition and aminoacylation, while Trp223, which in our analysis has a high correlation contact with A73, is essential to identify the discriminator base A73 for the selection of the correct tRNA molecule [61–63, 91]. Lys619 and Met620 are part of the conserved 619-KMSKS loop in the catalytic site, and make direct contacts with the ligand [60]. The loop shows great similarity to other conserved sequences found in ATP binding proteins, such as ATPsynthases, helicases and active transport pumps, and was postulated to help the *activation of the amino acid* by coordinating the *γ*-phosphate of ATP [93–96]. Since the initial state of the LeuRS-tRNA complex has an activated amino acid in its active site, the previously observed activation-related function of the 619-KMSKS motif was not relevant for the simulations carried out in the present study.

Interestingly, even though the whole KMSKS motif is conserved, previous studies that highlighted its importance focused on the -SKS end of the loop. The first lysine was shown to be essential and highly conserved, but the second lysine showed strong interactions with the pre-activation ATP substrate. Here, the initial two resides Lys619 and Met620 showed significant correlation with the adenylate, suggesting a role in stabilizing the ligand for the aminoacylation reaction. Moreover, by analyzing the betweeness measurements of the network, we observe that the edge with highest betweeness in the active site is the once connecting Lys619 to the tRNA base G71 (Fig. 6). This observation was consistent in all replicas of the system, suggesting a new information relay between active site and the C-terminal editing domain.

**Figure 6.**
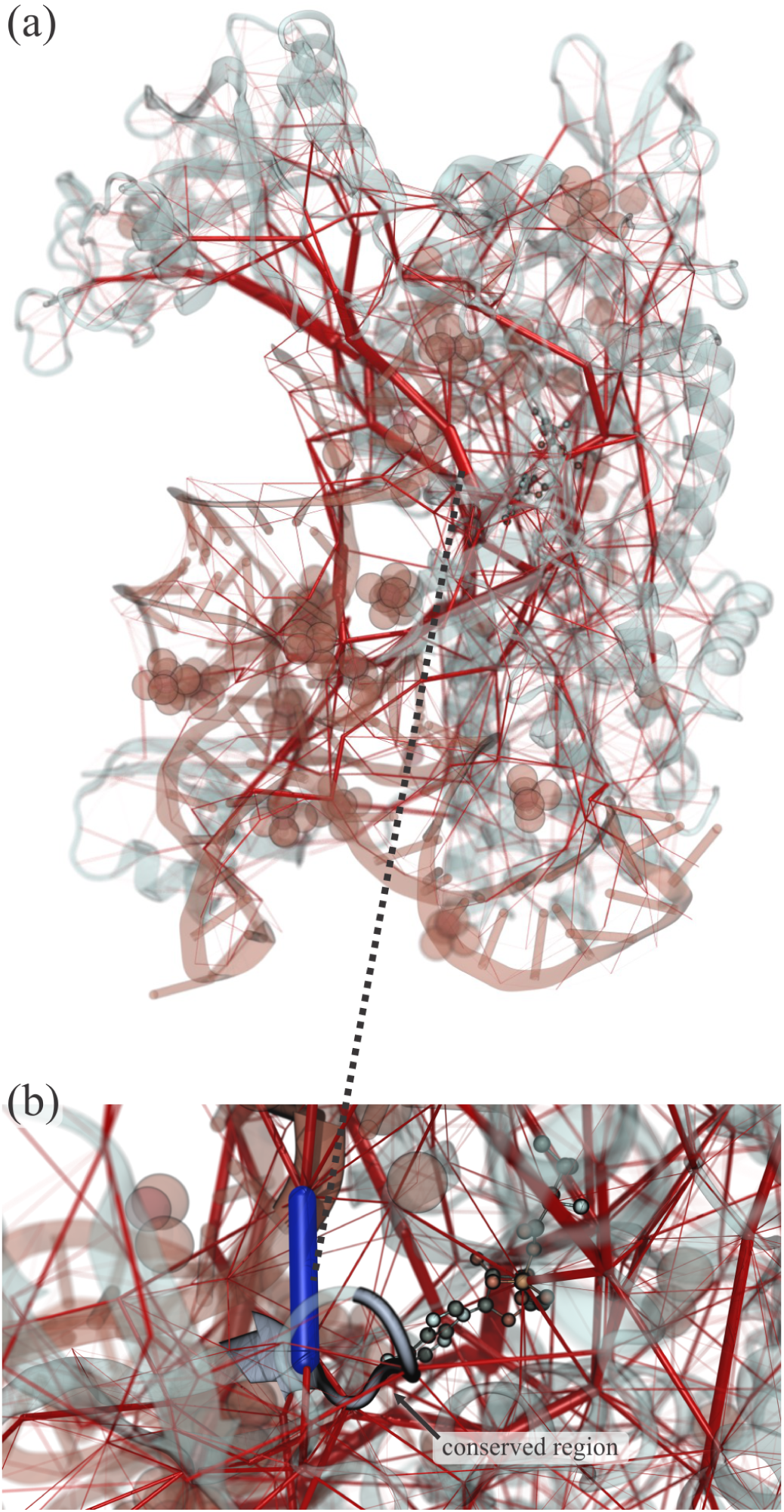
Rendering of LeuRS highlighting edge betweenness. (a) The transparent rendering of the protein and nucleic acid show the edges with high betweenness values (thick red tubes), connecting the catalytic region to the editing domain of LeuRS. (b) Zoom in the catalytic site showing the conserved 619-KMSKS loop in the catalytic site (in cartoon representation). The blue edge connects the protein from Lys619 to the tRNA, and is also the edge with highest betweenness value stemming from the active site. (Multimedia view)

Other contacts are also prominent in both systems, although no experimental evidence could be found to suggest a biochemical rationale for their role in complex stabilization. Examples are the Ala727-A22 and -C23 contacts, Asn734-C23 and -A24, Gly665-A24 and -C25 contacts, and Phe648-G12, -C23, and -A24 contacts. Bases C23 and A24 compose the core region of the tRNA, making them essential in keeping the overall structure of the tRNA. Also, Arg668 interacts with bases C25, A26, A38 and U39, making stabilizing contacts for the anticodon arm.

#### 4.2.4 Points of highest betweenness in the LeuRS

Perhaps a less frequently employed feature of dynamic networks analysis of biomolecules is the investigation of betweenness centrality. This feature can be defined as a measure of how important a network node is for communication within a biomolecule. For instance, how important an amino acid residue is in maintaining a protein’s activity. The betweenness is equal to the number of shortest paths from all vertices to all others that pass through that node. For instance, in a protein, amino acid residues in that have a high betweenness tend to be important for controlling inter-domain communication in that protein. Betweenness centrality can therefore be defined as a measure of centrality in a network based on shortest paths. For every pair of vertices in a network, there exists at least one shortest path between the vertices such that the sum of the generalized correlation coefficient of the edges is minimized. The betweenness centrality for each vertex is the number of these shortest paths that pass through the vertex.

Calculating the betweenness centrality of all the vertices in a network involves calculating the shortest paths between all pairs of vertices on that network, which for biomolecular systems is typically done with a modified Floyd–Warshall algorithm [43, 44]. The modifications allows this algorithm to find not only one but all shortest paths between any pair of nodes. In Fig. 6 we show the betweenness centrality of the LeuRS complex. The image depicts all the possible allosteric communications within the complex, showing what are the main communication hubs.

### 4.3 Respiratory Complex I

In the past four decades we have witnessed MD simulations evolving into a ‘computational microscope’. In the dawn of exascale-computing, high-performance MD software are enabling the investigation of large and complex systems, with many millions of atoms [2, 3]. These large-scale MD simulations are key to the understanding of the fundaments of life. But how do we analyze the complex mechanisms of communications within such large complexes. In this section we used the respiratory complex I, a transmembrane 2.5 M atoms system, to show a couple of interesting features that can be investigated using our generalized network analysis software.

#### 4.3.1 Identifying lipid-protein partners

Since a plethora of biochemical phenomena takes place within lipid membranes and their interface with water, investigating the interaction between lipids and other biomolecules is of high interest [97, 98]. For instance, G-Protein Couple Receptors are one of the most targeted protein classes for drug discovery, and they exist mainly as membrane-embedded proteins [99–101]. For these reasons, the lipid membrane has been investigated in many MD studies, both using a classical mechanics approach [102] or a hybrid QM/MM [103]. Having a large transmembrane segment, the respiratory complex I is a great example of a system where lipids might play a big role in regulating the protein’s activity. Beyond interactions protein subunits and individual lipids, the lateral forces offered by the biological membranes can also affect the dynamics of the protein complex. In fact, many transmembrane proteins are known to be mechanosensitive [104].

To investigate how the lipids in the membrane would interact with the respiratory complex I, we used a new feature of the Network Analysis tool that automatically searches for solvent or lipid residues that are stably bound to the biomolecular complex being studied. Therefore, we have assigned both the water and the lipid molecules that tightly interact with Complex I as part of the network. Here, as a simple test, we defined only one node per lipid, located in the phosphate group. Our network analysis revealed that, although not among the highest correlated movements (depicted as the red tubes in Fig. 7a), many lipids could be considered part of the respiratory complex I as they remain in constant contact with the protein throughout sections of the simulation. In Fig. 7a one can identify several regions of the respiratory complex I that are highly correlated, particularly in the regions exposed to the water solvent. The lipids that were considered part of the network are highlighted in Fig. 7b. It is noteworthy that these lipids are concentrated in specific regions of the protein complex. We should, however, emphasize that these are relatively short MD simulations (100 ns long), that serve as tests for our software. A better understanding of these protein-lipids interaction is still an open question that would find necessary many replicates of longer MD simulations, and associated with experimental evidences.

**Figure 7.**
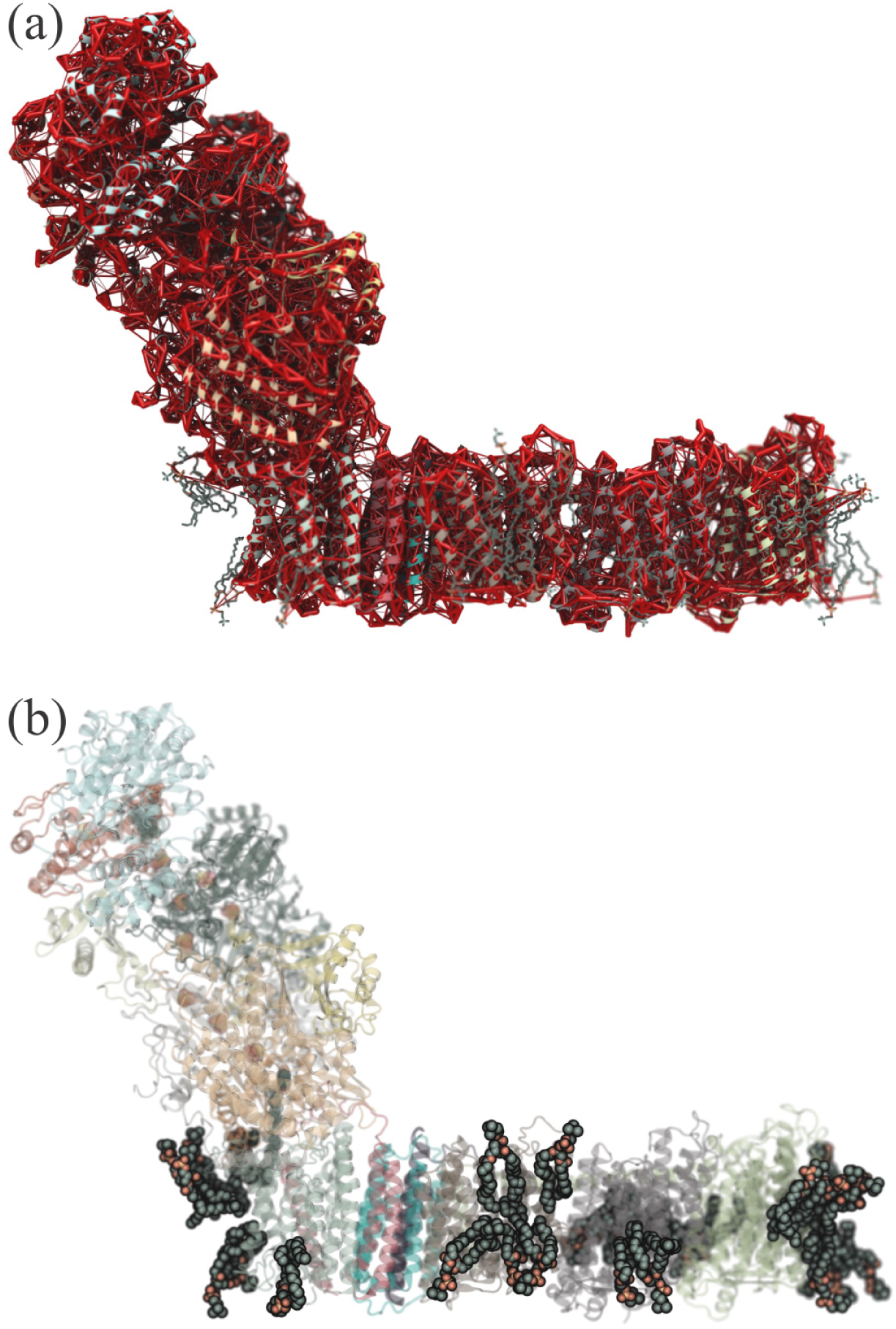
The interaction of lipids with the respiratory complex I. (a) Full network revealing the most correlated regions of the complex. Edge thickness represents their generalized correlation coefficient. (b) Rendering highlighting the lipids that were found to be stably bound and highly correlated to the protein complex. The protein representation is colored by communities.

#### 4.3.2 Allosteric communication and electron transfer

In the respiratory complex I, the peripheral-arm, is responsible for removing two electrons from an NADH, which are then transferred to a quinone through a bridge formed by a flavin and eight iron-sulfur complexes. An additional iron-sulfur complex is known to be off the main redox pathway [69]. The elaborate mechanism of the respiratory complex I leads to the addition of two electrons and two protons to the quinone, converting it to a quinol. This conversion is then known to induce the activity of four proton pumps located in the transmembrane-arm of the respiratory complex I. This chemo-mechanical coupling connect processes that span only picoseconds to conformational transitions that happen in the millisecond timescale [69]. Therefore, the allosteric pathway connecting the flavin and the quinone in the respiratory complex I structure is of key interest. For a low-barrier transfer of the electrons through the peripheral-arm, a stable path is need. Using our generalized network analysis tool, we can calculate these allosteric pathways with high correlation sensitivity.

The dynamical network models of allostery can identify optimal and suboptimal allosteric pathways. The statistical distribution of these pathways are useful for locating accessible residues that can work as allosteric regulators of important drug targets [19]. Here, the Floyd-Warshall algorithm [43, 44], which uses the correlations as weights to calculate network distances and shortest distances, was employed to find the optimal and suboptimal pathways connecting the flavin and the quinone in the respiratory complex I structure. To identify the suboptimal paths, this algorithm searches for the optimal (shortest) path, with all other paths deemed suboptimal if they fall within an acceptable deviation from the optimal path.

In Fig. 8 one can observe not only what are the suboptimal pathways, but how they evolve over time. One of the features of the generalized network analysis software is that the user can easily breakdown the trajectories into windows. These windows represent the evolution of the system over time, as the correlation is only calculated within that window. Such feature will assist our users in identifying when a pathway is more stable for a QM/MM calculation, or how force-propagation pathways evolve in a single-molecule force spectroscopy experiment. It is noteworthy that the suboptimal pathways frequently present extremely degenerated signal along the optimal pathway. That is clear in our Fig. 8, where in some parts of the system a very degenerate signal is found. However, our results clearly indicate how the pathways more or less follow the chain of iron-sulfur complexes.

**Figure 8.**
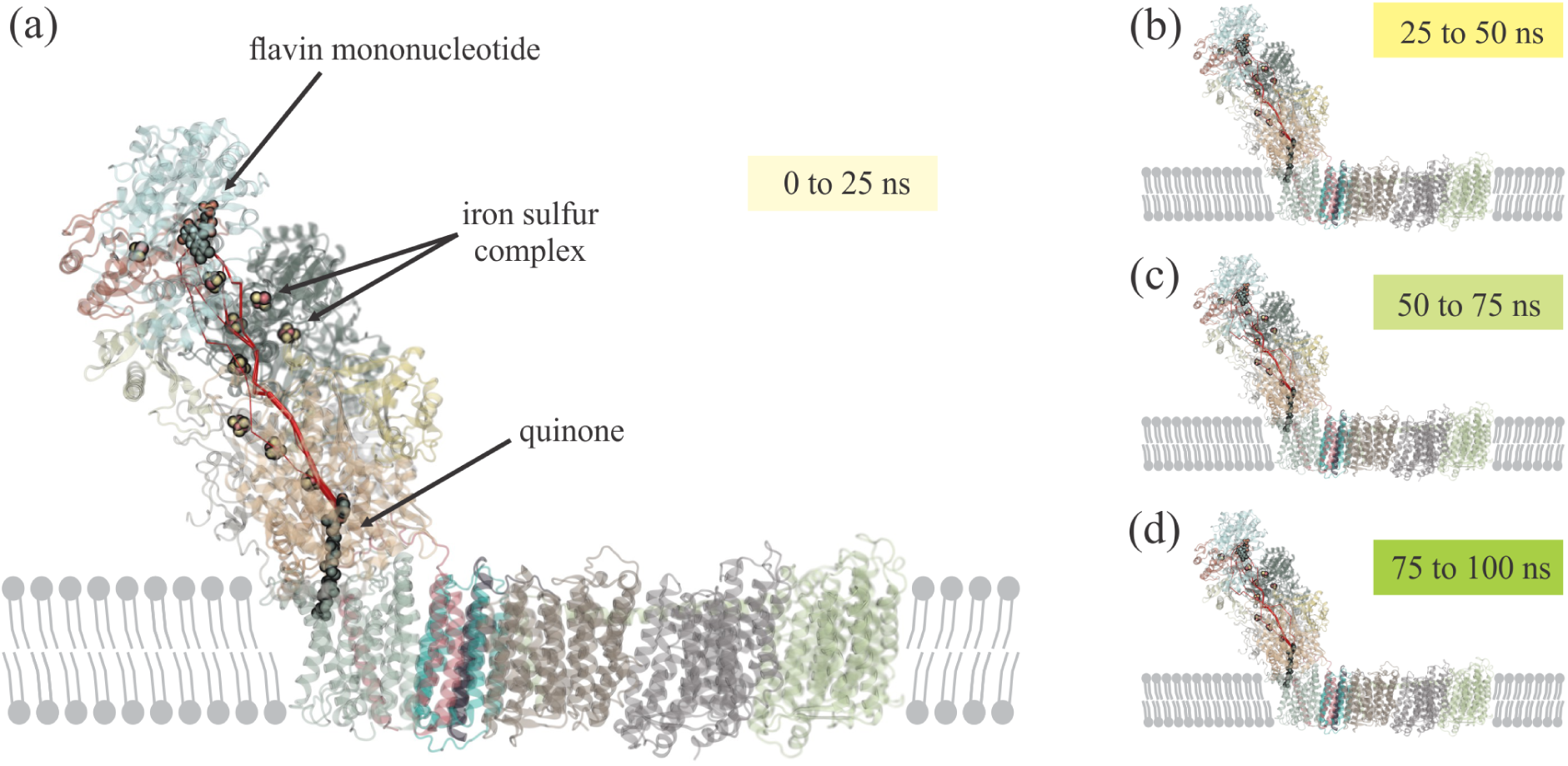
Rendering of the time evolution of the suboptimal paths connecting the flavin mononucleotide and the quinone in the respiratory complex I. (a) First simulation window: from 0 ns to 25 ns. (b) Second simulation window: from 25 ns to 50 ns. (c) Third simulation window: from 50 ns to 75 ns. (b) Forth simulation window: from 75 ns to 100 ns. The image highlights the iron sulfur centers that serve to carry electrons from the NADH dehydration site to the quinone. All renderings were produced with VMD and our new Network Viewer 2.0 GUI.

## 5 Concluding Remarks

The increasing use of MD simulations for ever-larger systems challenges not only the MD engines, but also the analysis tools used to investigate MD trajectories. Another common problem created by the large amount of data generated is how to filter such data in trajectory analysis. A popular strategy has been to use artificial intelligence algorithms to search for patterns in these trajectories. Another important strategy is to look into how distant parts of a molecular system can “talk” to one another. For the latter, dynamical network analysis theory is perhaps the most appropriate way to investigate these large set of trajectories. Dynamical network analysis tools are typically optimized to work with large-scale networks that are formed by nodes and the links between these nodes, commonly called edges. In a molecular system these nodes can represent one molecule, for instance an amino acid residue, a group of molecules, or even a small group of atoms within a molecule. The links between these nodes are typically assigned based on proximity of the nodes and the correlation in the motion between these nodes.

The most common approach to calculate correlation in MD trajectories is to employ Pearson correlation coefficients [105]. However, despite being inexpensive to calculate, Pearson coefficients do not account for non-linear contributions to correlation [48]. Generalized correlation coefficients can account for non-linear contributions, but are more computationally expensive to calculate. Here, we have presented a new implementation of network analysis that takes advantage of the sparsity of the network correlation matrix. Since each node is only ever in contact with few other nodes, the expensive generalized correlation calculation could be transitioned from an *N* ^2^ problem to a linear calculation. We have tested our new software in three different systems, with different levels of complexity.

With enhanced efficiency and ease-of-use, the generalized network analysis software can be applied to study transient information transfer between biomolecules in crowded environments, such as a cell’s cytoplasm. As large scale simulations are currently being published or under way [3], using all-atom and coarse grained representations, reaching dozens of millions of atoms, no tool was able to produce generalized correlation networks for them. Due to the new way of calculating the network correlation matrix, our software is able to cut the analysis time from 12 hours to just 3 minutes for a system of the size of a ribosome, or from 4 days to 8 minutes for the whole HIV capsid. At the same time, multiple replicas can be analyzed in parallel.

As a test case, we have used replicas representing independent simulations of the same system (LeuRS), but these replicas could be used to describe a mechanical changes, large scale motions achieved through enhanced sampling techniques, or millisecond long simulations of biomolecules. We have shown that network analysis can recover essential contacts in the protein:tRNA interface of the LeuRS complex. Moreover, the updated implementation and interface make use of latest technologies to provide fast analysis and informative results, as well as an interactive environment that allows the exploration of features particular to each individual system. In particular, the linear scalability in correlation calculation afforded by the current implementation allows for large systems to be tackled, without compromising precision of correlation values or coverage of the macro-molecular system.

We have also provided a comprehensive tutorial to investigate the OMP-decarboxylase, which will allow our users to quickly adapt the generalized network analysis to their analysis routine. Additionally, we have looked into a third system, namely the respiratory complex I, showing how membrane’s lipids become effective part of a protein network. The same approach can be used to investigate which lipids are modulating other transmembrane proteins, or to identify lipids that have a highly correlated movement in a membrane, as in a lipid raft.

In summary, we have developed a generalized network analysis software that allows for the investigation of MD trajectories of large biomolecular complexes. Our software, which is implemented as a python package, provides a pipeline for investigating many network properties, such as community and allosteric pathway analysis. A new VMD script provides an easy-to-use menu for production of high-quality renderings of the network maps, as presented throughout the manuscript.

This work was supported by the National Institutes of Health (NIH) grant P41-GM104601, and by the National Science Foundation (NSF) grant MCB-1616590. Molecular dynamics simulations made use of XK-nodes of NCSA Blue Waters supercomputer, which are Nvidia GPU-accelerated. The state of Illinois and the National Science Foundation (awards OCI-0725070 and ACI-1238993) support Blue Waters sustained-petascale computing project. Cesar de la Fuente-Nunez holds a Presidential Professorship at the University of Pennsylvania, is a recipient of the Langer Prize by the AIChE Foundation and acknowledges funding from the Institute for Diabetes, Obesity, and Metabolism and the Penn Mental Health AIDS Research Center of the University of Pennsylvania.

## Supporting information

(Multimedia view)

## Supplementary Information

The necessary information to reproduce our results is presented in the form of a tutorial as supplementary information. Simulation data will be provided by authors upon reasonable request. All Python and TCL code necessary for analysis and plots are provided as supplementary material.

## Publication Note

The following article has been submitted to The Journal of Chemical Physics. After it is published, it will be found at *https* : *//aip*.*scitation*.*org/journal/jcp*.

