## Supplementary material for "Generalized correlation-based dynamical network analysis: a new high-performance approach for identifying allosteric communications in molecular dynamics trajectories": environment.yml

```
name: DyNetAnTutorial

channels:
  - conda-forge
  - defaults
  
dependencies:
    - python>=3.8.*
    - pip>=20.*
    - colorama
    - h5py
    - jupyter
    - ipywidgets
    - mdanalysis
    - networkx
    - nglview
    - numba
    - numpy
    - pandas
    - pympler
    - python-louvain
    - scipy
    - tzlocal
    - rpy2
    - r-data.table 
    - r-ggplot2
    - r-ggrepel
    - r-gdata
    - r-rcolorbrewer
    - r-colorramps
    - r-rpref
    - pip:
        - dynetan
```
