## Supplementary material for "Generalized correlation-based dynamical network analysis: a new high-performance approach for identifying allosteric communications in molecular dynamics trajectories": README.md

# Dynamic Network Analysis - Tutorial
A tutorial for the application of Dynamical Network Analysis using Generalized Correlations.

## Installing Requirements

There are two main ways to instll all dependencies to execute this tutorial: The first is by creating a Conda environment, the second is by manually installing system packages, and Python and R packages. We provide examples for both method below.

Building a Conda Environment
----------------

The easyest way to install all requirements for the execution of this tutorial is to create a Conda environment. Inside the folder *CondaEnv* you will find a recipe for an environment with all required Python and R packages. 
To create the environment using a terminal (a command line interface), simply go to the *CondaEnv* folder and run the following command:

    conda env create -f environment.yml

Custom Installation
----------------

**System Packages**

For this tutorial, you will need an R installation (usually provided by system packages `R-core` and `R-core-devel`) and system packages for the "gdata" package (usually provided by system packages `perl-Compress-Raw-Zlib` and `perl-Spreadsheet-XLSX`). In Fedora 32, all can be installed with the following command:

    dnf install perl-Compress-Raw-Zlib perl-Spreadsheet-XLSX R-core-devel

**Python Packages**

To run the tutorial Jupyter Notebooks, you will need the `dynetan` package and `rpy2`. The following commands will install all required python packages, assuming you have met all system requirements.

    pip install dynetan
    pip install rpy2

To activate notebook widgets (from the ipywidgets package) and visualization (from the nglview package), you may need to execute the following commands:

    jupyter nbextension enable --py widgetsnbextension
    jupyter-nbextension enable nglview --py --user

**R Packages**

To install R packages, execute the following commands in your R terminal:

    list.of.packages <- c("data.table", "ggplot2", "ggrepel", "gdata", "RColorBrewer", "colorRamps", "rPref")

    new.packages <- list.of.packages[!(list.of.packages %in% installed.packages()[,"Package"])]

    if(length(new.packages)) install.packages(new.packages, repos='http://cran.wustl.edu/', dependencies = TRUE)
