## Supplementary material for "Generalized correlation-based dynamical network analysis: a new high-performance approach for identifying allosteric communications in molecular dynamics trajectories": Tutorial-Step_1-ProcessTrajectory.ipynb

```
# %% [markdown]
# Generalized Network Analysis Tutorial - Step 1

The Network Analysis Tutorial is part of the work entitled **Generalized correlation-based dynamical network analysis: a new high-performance approach for identifying allosteric communications in molecular dynamics trajectories**, by Marcelo C. R. Melo, Rafael C. Bernardi, Cesar de la Fuente-Nunez, and Zaida Luthey-Schulten. For more information see http://faculty.scs.illinois.edu/schulten/. 

In this tutorial, we will use the Dynamic Network Analysis python package to explore the interactions between the OMP decarboxylase enzyme and its substrate, identifying clusters of amino acid residues that form domains, and active site residues that are important for binding and enzymatic activity.

The tutorial is divided in two jupyter notebooks. In this notebook, **Step 1**, we will analyze the MD trajectory for the OMP decarboxylase system and generate the network data used for analysis. 

The trajectory files have approximately 500MB in size, and must be downloaded [from this link](http://www.ks.uiuc.edu/~rcbernardi/NetworkAnalysis/DynamicNetworkAnalysis_MDdata.tar.gz) and placed in the *TutorialData* folder.

In the accompanying notebook, **Step 2**, we will load the data generated in the Step 1, and create analysis plots and visualizations of the network. 

In **Step 3**, which is outside of the jupyter notebook environment, we will produce high-quality renderings of the network using the popular visualization software VMD.


**Important 1:**

If you are not familiar with Jupyter Notebooks, please take some time to watch a video tutorial, such as this:

https://www.youtube.com/watch?v=HW29067qVWk

If you are not familiar with VMD, you can find tutorials on how to use VMD here:

https://www.ks.uiuc.edu/Training/Tutorials/


**Important 2:**

Here, we use a language that is common among computational structural biophysicists. For instance:

The word **residue** is typically used to represent both an amino acid residue, or any other small molecule in the system, such as a ligand.

The word **segment** and also **segID** is typically used to represent a polymer chain, such as a protein, or a group of molecules in a lipid bilayer. Protein complexes would therefore be formed by multiple segments.


# %% [code]
# Load the python package
import os
import dynetan
from dynetan.toolkit import *
from dynetan.viz import *
from dynetan.proctraj import *
from dynetan.gencor import *
from dynetan.contact import *

# Create the object that processes MD trajectories.
dnap = DNAproc()

# %% [markdown]
# Node Groups

In Network Analysis, each residue is represented by one or more "nodes", serving as proxies for groups of atoms form the original residue (Figure 1). This approach lowers computational cost and noise. 

![](./toolkit/figures/nodeAtomGroup.png)
__Figure 1:__ (a) and (b) represent a chain of amino acids and a nuceotide, respectively. (c) and (d) represent the same molecules but highlighting the difference between a node (alpha carbon in amino acids, and nitrogen and phosphate in nucleotides), and the group of atoms that the node represents.

# System info:
To get started, we need some basic information about the system:
- Work directory where we will find relevant files.
- Names of PSF and DCD files.
- segid's for the regions that will be used for network analysis.
- Name of the solvent residue(s) [default: TIP3]
- Number of windows into which the trajectory will be split. [default: 4]
- Name of atom(s) that will represent node(s) in each residue.
- Names of atoms in node groups.

**You can can find a complete list of supported trajectory formats [here](https://www.mdanalysis.org/docs/documentation_pages/coordinates/init.html#supported-coordinate-formats):**


# %% [code]
# Path where input files will searched and results be written.
workDir = "./TutorialData/"

# PSF file name
psfFile = os.path.join(workDir, "decarboxylase.0.psf")

# DCD file name
dcdFiles = [os.path.join(workDir, "decarboxylase.1.dcd")]
# dcdFiles = [os.path.join(workDir, "decarboxylase.1.short.dcd")]

ligandSegID = "OMP"

# Segment IDs for regions that will be studied.
segIDs = ["OMP","ENZY"]

# Residue name for solvent molecule(s)
h2oName = ["TIP3"]

# Number of windows created from full simulation.
numWinds = 4

# Sampled frames per window
numSampledFrames = 10

# Number of sampled frames for automatic selection of solvent and ions.
# numAutoFrames = numSampledFrames*numWinds

# Network Analysis will make one node per protein residue (in the alpha carbon)
# For other residues, the user must specify atom(s) that will represent a node.
customResNodes = {}
customResNodes["TIP3"] = ["OH2"]
customResNodes["OMP"] = ["N1","P"]

# We also need to know the heavy atoms that compose each node group.

usrNodeGroups = {}

usrNodeGroups["TIP3"] = {}
usrNodeGroups["TIP3"]["OH2"] = set("OH2 H1 H2".split())

usrNodeGroups["OMP"] = {}
usrNodeGroups["OMP"]["N1"] = set("N1 C2 O2 N3 C4 O4 C5 C6 C7 OA OB".split())
usrNodeGroups["OMP"]["P"] = set("P OP1 OP2 OP3 O5' C5' C4' O4' C1' C3' C2' O2' O3'".split())

#################################
### Extra configuration

# Cutoff for contact map (In Angstroms)
cutoffDist = 4.5

# Minimum contact persistance (In ratio of total trajectory frames)
contactPersistence = 0.75

#################################
### Load info to object

dnap.setNumWinds(numWinds)
dnap.setNumSampledFrames(numSampledFrames)
dnap.setCutoffDist(cutoffDist)
dnap.setContactPersistence(contactPersistence)
dnap.seth2oName(h2oName)
dnap.setSegIDs(segIDs)

dnap.setCustomResNodes(customResNodes)
dnap.setUsrNodeGroups(usrNodeGroups)

# %% [markdown]
# Load the trajectory

Our Generalized Network Analysis leverages the MDAnalysis package to create a *universe* that contains all the trajectory and system information.

# %% [code]
dnap.loadSystem(psfFile,dcdFiles)

# %% [code]
# We can access the trajectory data directly.
dnap.getU().trajectory

# %% [markdown]
### Checks segments and residue names

This is important to know if there are residues in the structure that we didn't know of, and need to be addresssed so that network analysis can create nodes in all selected residues.

# %% [code]
dnap.checkSystem()

# %% [code]
# More checks?
#
# You can add more checks here if your system has non-standard residues that need special attention.

# %% [markdown]
### Automatically identify crystallographic waters and ions.

In this section we identify all residues which will be checked for connectivity with selected segments.

- First, we define if all solvent molecules will be checked, or if just ions, ligands, lipids, and other molecules will be checked.

- Second, we sample a small set of frames from the trajectory to select likely residues, then we check all trajectory to see if they are closer than the cutoff distance for at least x% of the simulation (where x is the "contact persistence" fraction of the trajectory).

- Third, we load the trajectory of the relevant atoms to memory to spped up the analysis.

# %% [code]
dnap.selectSystem(withSolvent=True)

# %% [markdown]
### Prepare network representation of the system

Here we check that we know how to treat all types of residues in the final selection. Every residue will generate one or more nodes in the final network. Then we store the groups of atoms that define each node.

# %% [code]
dnap.prepareNetwork()

# %% [markdown]
## Align the trajectory based on selected segments

We align the trajectory to its first frame using heavy atoms (non-hydrogen) from the selected segments. In the process, we also transfer the trajectory to the computer memory, so that future analysis and manipulations are completed faster.

# %% [code]
# If your system is too large, you can turn off the "in memory" option, at a cost for performance.
dnap.alignTraj(inMemory=True)

# %% [markdown]
### Select residues that are closer than 4.5A for more than 75% of simulation

<img align="right" src="./toolkit/figures/nodeContactDistances.png" alt="drawing" width="400px"/>

Creates an N-by-N matrix for all N nodes in the selected region, and automatically selected nodes (ions, solvent).

The following cell defines efficient functions to run the analysis and create a contact matrix. We leverage both MDAnalysis parallel contact detection tools, as well as accelerated Numba and Cython function. After creating the contact matrix, we remove any automatically selected nodes that have insuficient persistance, and filter the contacts by (optionally) removing contacts between nodes in consecutive residues in a protein or nucleic chain.

In the example to the right, the distances between atoms indicates that in that frame of the simulation, the node representing the water molecule would make a contact with the phosphate node (representing the ribose ring and phosphate), but not with the nitrogen node (representing the nucleic base).


# %% [markdown]
**Attention** For every time you start this Jupyter Notebook, the first time you execute this function may take significanlty longer (several seconds) to start. This is because we use *Cython* and *Numba* to compile functions "on-demand", and a new compilation may be necessary after the notebook is re-started.

# %% [code]
# To speed-up the contact matrix calculation, a larger stride can be selected, at a cost for precision.
dnap.findContacts(stride=1)

# %% [markdown]
### Removing contacts between nodes in the same residue.

The following function guarantees that there will be no "self-contacts" (contacts between a node and itself), and gives you the opportunity to remove contacts between nodes in consecutive residues (such as sequential amino acids in the same chain, removing back-bone interactions). 

The function also removes nodes that are never in contact with any other node in the system (such as the ends of flexible chains, or residues in flexible loops). This will automatically update the MDanalysis universe and related network informatio, such as number of nodes and atom-to-node mappings.

# %% [code]
dnap.filterContacts(notSameRes=True, notConsecutiveRes=False, removeIsolatedNodes=True)

# %% [markdown]
## Calculate Generalized Correlation with Python/Numba

# %% [code]
# We can calculate generalized correlaions in parallel using Python's multiprocessing package.
dnap.calcCor(ncores=4)

# %% [markdown]
## Calculate cartesian distances between all nodes in the selected system.

Here, we will calculate the **shortest** distance between atoms in all pairs of nodes. It is similar to the contact matrix calculation, but we check all distances and keep the shortest one to use in our analysis.

# %% [code]
# We can leverage MDanalysis parallelization options with backend="serial" or backend="openmp".
# For very small systems, the serial can be faster!
dnap.calcCartesian(backend="serial")

# %% [markdown]
# Network Calculations
Create a graph from our correlation matrix. Different properties are calculated:

*Density* maesures how connected the graph is compared to how connected it *could* be. It is the ratio between edges in the graph over all possible edges between all pairs of nodes.

*Transitivity* maesures the triadic closure, comparing present triangles to possible triangles. In a triangle, if A is connected to B, and B connected to C, then A is connected to C.

*Degree* measures the number of connections a node has.

(Reference)[1]

[1]:https://programminghistorian.org/en/lessons/exploring-and-analyzing-network-data-with-python#advanced-networkx-community-detection-with-modularity

# %% [code]
dnap.calcGraphInfo()

# %% [code]
# Basic information of the network as interpreted as a graph.
print( nx.info(dnap.nxGraphs[0]))

# %% [code]
# Both density and transitivity are scaled from 0 to 1
for win in range(dnap.numWinds):
    print("----- Window {} -----".format(win))
    print("Density:", round( nx.density(dnap.nxGraphs[win]), 4) )
    print("Transitivity:", round( nx.transitivity(dnap.nxGraphs[win]), 4) )
    print()

# %% [code]
from operator import itemgetter

# We can check the nodes that have the most connections in each window.
for win in range(dnap.numWinds):
    print("----- Window {} -----".format(win))
    
    sorted_degree = sorted(dnap.getDegreeDict(win).items(), key=itemgetter(1), reverse=True)
    
    print("Top 10 nodes by degree: [node --> degree : selection]")
    for n,d in sorted_degree[:5]:
        print("{0:>4} --> {1:>2} : {2}".format(n, d, getSelFromNode(n, dnap.nodesAtmSel)))
    
    print()

# %% [markdown]
## Calculate optimal paths
We choose the Floyd Warshall algorithm[1]. This uses the **correlations as weights** to calculate network distances and shortest distances.

[1]:https://networkx.github.io/documentation/stable/reference/algorithms/generated/networkx.algorithms.shortest_paths.dense.floyd_warshall.html?highlight=warshall#networkx.algorithms.shortest_paths.dense.floyd_warshall

# %% [code]
dnap.calcOptPaths(ncores=4)

# %% [markdown]
## Calculate betweenness

We calculate both betweenness centrality[1] for edges and eigenvector centrality[2] for nodes.

[1]:https://networkx.github.io/documentation/stable/reference/algorithms/generated/networkx.algorithms.centrality.edge_betweenness_centrality.html?highlight=betweenness#networkx.algorithms.centrality.edge_betweenness_centrality
[2]:https://networkx.github.io/documentation/networkx-1.10/reference/generated/networkx.algorithms.centrality.eigenvector_centrality.html

# %% [code]
dnap.calcBetween(ncores=4)

# %% [code]
from itertools import islice

# Pairs of nodes with highest Betweeness values, compared to their correlation values (in Window 0)
for k,v in islice(dnap.btws[0].items(),5):
    print("Nodes {} have betweenes {} and correlation {}.".format(k, 
                                                                  round(v,3), 
                                                                  round(dnap.corrMatAll[0, k[0], k[1]], 3) ) )

# %% [markdown]
Turn to node centrality instead of edge centrality:

# %% [code]
dnap.calcEigenCentral()

# %% [markdown]
### Calculate communities


# %% [markdown]
Using **Louvain heuristices** is feasible. 
This method also maximizes the modularity of the network.

http://iopscience.iop.org/article/10.1088/1742-5468/2008/10/P10008/meta

# %% [code]
dnap.calcCommunities()

# %% [code]
# Sort communities based on number of nodes
for comIndx in dnap.nodesComm[0]["commOrderSize"]:
    print("Modularity Class {0:>2}: {1:>3} nodes.".format(comIndx, len(dnap.nodesComm[0]["commNodes"][comIndx])))

# %% [code]
# Sort communities based on the node with highest eigenvector centrality
for comIndx in dnap.nodesComm[0]["commOrderEigenCentr"]:
    print("Modularity Class {0} ({1} nodes) Sorted by Eigenvector Centrality:".format(
                                                                    comIndx, 
                                                                len(dnap.nodesComm[0]["commNodes"][comIndx])))
    for node in dnap.nodesComm[0]["commNodes"][comIndx][:5]:
        print("Name: {0:>4} | Degree: {1:>2} | Eigenvector Centrality: {2}".format(
            node, dnap.nxGraphs[win].nodes[node]['degree'], dnap.nxGraphs[win].nodes[node]['eigenvector']))
    print()

# %% [markdown]
# Process Interface Residues

We now find all nodes that are close to both selections chosen by the user. That may include amino acids in the interface, as well as ligands, waters and ions.

# %% [code]
dnap.interfaceAnalysis(selAstr="segid ENZY", selBstr="segid OMP")

# %% [markdown]
## Save The Data

# %% [code]
pathToData = "./TutorialResults/"

fileNameRoot = "dnaData"

fullPathRoot = os.path.join(pathToData, fileNameRoot)

dnap.saveData(fullPathRoot)

# %% [markdown]
## Save reduced trajectory for visualization

# %% [code]
# This function will save a reduced DCD trajectory with the heavy atoms used for network analysis
# A smaller trajectory can be created by choosing a "stride" that sub-samples the original trajectory.
# This function will also produce a PDB file so that information on atoms and residues can be loaded to
#    visualization software such as VMD.

dcdstride = 1

print("We will save {} heavy atoms and {} frames.".format(dnap.workU.atoms.n_atoms, 
                                                          len(dnap.workU.trajectory[::dcdstride]) ))

# %% [code]
dnap.saveReducedTraj(fullPathRoot, stride = dcdstride)

# %% [markdown]
MDAnalysis may print warnings regarding missing data fields, such as altLocs, icodes, occupancies, or tempfactor, which provide information commonly found in PDB files.
The warnings are for your information and in the context of this tutorial they are expected and do not indicate a problem.

# %% [markdown]
# Analysis

The we have finished processing the trajectory and storing all related data. We can now move on to analysis of the network properties calculated here.

**All analysis code was placed in a second tutorial notebook for clarity.**


# ---- The End ----
```
