## Supplementary material for "Generalized correlation-based dynamical network analysis: a new high-performance approach for identifying allosteric communications in molecular dynamics trajectories": Tutorial-Step_2-AnalysisAndPlots.ipynb

```
# %% [markdown]
# Generalized Network Analysis Tutorial - Step 2

In the second part of our generalized network analysis tutorial, the user is presented with another jupyter notebook where all the correlation maps calculated in the first part can be translated into molecular visualizations or plots. Here we provide an opportunity for users more comfortable with python programming to tailor the analysis to their specific scientific questions.

The Network Analysis Tutorial is part of the work entitled **Generalized correlation-based dynamical network analysis: a new high-performance approach for identifying allosteric communications in molecular dynamics trajectories**, by Marcelo C. R. Melo, Rafael C. Bernardi, Cesar de la Fuente-Nunez, and Zaida Luthey-Schulten. For more information see http://faculty.scs.illinois.edu/schulten/. 

# %% [code]
# Load the python package
import os
from dynetan.toolkit import *
from dynetan.viz import *
from dynetan.proctraj import *
from dynetan.gencor import *
from dynetan.contact import *
from dynetan.datastorage import *

#from numpy.linalg import norm
from itertools import islice
from scipy import stats

import pandas as pd
import numpy as np
import scipy as sp

# For visualization
import nglview    as nv
import ipywidgets as widgets

# %% [code]
dnad = DNAdata()

# %% [code]
# Allows better plotting inside jupyter notebooks
%matplotlib notebook

# %% [code]
%load_ext rpy2.ipython

# %% [code]
# Make sure all the following packages are installed in your R distribution

%R require(data.table)
%R require(ggplot2)
%R require(ggrepel)
%R require(gdata)
%R require(RColorBrewer)
%R require(colorRamps)
%R require(rPref)

# %% [markdown]
## Files and system definitions (same as in "ProcessTrajectory" notebook) 

Make sure that all the paths are the same as in the **Step 1** notebook. Also, you might want to have analysis resutls saved to a new location, to avoid mixing results of different systems. If you are copying this tutorial to adapt to your own system, be careful to select new paths. 

# %% [code]
dataDir = "./TutorialResults/"

# Path where results will be written (you may want plots and data files in a new location)
workDir = "./TutorialResults/AnalysisResults"

fileNameRoot = "dnaData"
fullPathRoot = os.path.join(dataDir, fileNameRoot)

# Define the segID of the Ligand being studied.
ligandSegID = "OMP"

# %% [code]
# Creates the workdirectory, and Plots directory.
import os
if not os.path.exists(workDir):
    os.makedirs(workDir)
    os.makedirs(os.path.join(workDir, "Plots"))

# %% [markdown]
## Load the Data

# %% [code]
dnad.loadFromFile(fullPathRoot)

# %% [markdown]
Load reduced trajectory, in case this notebook did not load ALL trajectories.

# %% [code]
dcdVizFile = fullPathRoot + "_reducedTraj.dcd"
pdbVizFile = fullPathRoot + "_reducedTraj.pdb"

workUviz = mda.Universe(pdbVizFile, dcdVizFile)

# %% [markdown]
If we are loading the first trajectory of the notebook, we need to create the dnad.nodesAtmSel structure to make selections on the universe based on node indices. This is mostly used for visualizations.

# %% [code]
# We add this to the object for ease of access.
dnad.nodesAtmSel = workUviz.atoms[ dnad.nodesIxArray ]

# %% [code]
trgtNodes = getNodeFromSel("segid " + ligandSegID, dnad.nodesAtmSel, dnad.atomToNode)
trgtNode = getNodeFromSel("segid " + ligandSegID + " and name P", dnad.nodesAtmSel, dnad.atomToNode)
print(trgtNodes, trgtNode)

# %% [markdown]
# Load and Visualize Paths and Communities


# %% [markdown]
### The following cells will:

- Open an NGL View window displaying the molecular system. If the molecular system is not loaded check for possible errors in loading the python modules.
- Show the optimal path between two nodes.


# %% [code]
# http://proteinformatics.charite.de/ngl/doc/#User_manual/Usage/Selection_language

w = nv.show_mdanalysis(workUviz.select_atoms("all"))

w._remote_call("setSize", target="Widget", args=["800px", "600px"])
w.parameters = dict(theme='light')

w.clear_representations()
w.add_cartoon("protein")
w.add_hyperball(ligandSegID)
w.add_hyperball("not protein")

w

# %% [code]
# Select an arbitrary node in the enzyme
enzNode = getNodeFromSel("segid ENZY and resname GLU and resid 115", dnad.nodesAtmSel, dnad.atomToNode)

# Get the optimal path connecting the two nondes.

optpath = getPath( trgtNodes[1], enzNode[0], dnad.nodesAtmSel, dnad.preds)

viewPath(w, optpath, dnad.distsAll, dnad.maxDirectDist, dnad.nodesAtmSel)

# %% [code]
w.close()

# %% [markdown]
### The following cells will:
- Open an NGL View window displaying the system.
- Show the contacts with highst betweeness.

# %% [code]
# http://proteinformatics.charite.de/ngl/doc/#User_manual/Usage/Selection_language

w = nv.show_mdanalysis(workUviz.select_atoms("all"))

w._remote_call("setSize", target="Widget", args=["800px", "600px"])
w.parameters = dict(theme='light')

w.clear_representations()
w.add_cartoon("protein")
w.add_hyperball(ligandSegID)
w.add_hyperball("not protein")

w

# %% [code]
####
# Plot the 10 highest betweennesses (edges most used by optimal paths)
####

w.clear_representations()
w.add_cartoon("protein")
w.add_hyperball(ligandSegID)
w.add_hyperball("not protein")

window = 0
for k,v in islice(dnad.btws[window].items(),10):
    print(k,v, dnad.corrMatAll[0, k[0], k[1]])
    viewPath(w, getPath(k[0], k[1], dnad.nodesAtmSel, dnad.preds), 
             dnad.distsAll, dnad.maxDirectDist, dnad.nodesAtmSel, window)

# %% [code]
####
# Save the current visualization in a figue.
####

#w.download_image(filename='nglview_frame.png', factor=15, trim=True, antialias=True)

# %% [code]
####
# Close the current visualization.
####

w.close()

# %% [markdown]
## Analyze Communitites 
We keep the communities that have more than 1% of nodes in all windows. Then we group communities across replicas by largest intersection. This is needed because we have no guarantee that the same community will be assigned the same ID in different windows of the same simulation.

We fin ally rank the communities by modularity.

# %% [code]
import networkx.algorithms.community.quality as nxquality

# Creates a list of windows and order them according to graph modularity.
windModul = []
for window in range(dnad.numWinds):
    modul = nxquality.modularity(dnad.nxGraphs[window], 
                         [ set(nodesList) for nodesList in dnad.nodesComm[window]["commNodes"].values()])
    windModul.append((window, modul))
    
windModul.sort(key=lambda x:x[1], reverse=True)

# Keep the window with the highest modularity as a reference for community matching
refWindow = windModul[0][0]

for wind, mod in windModul[:5]:
    print( "Window {} has modularity {:1.4f}.".format(wind, mod) )

# %% [code]
cutoff = max(10, np.ceil(0.01*dnad.numNodes))

def matchComm(mCommID, mWindow, refWindow, dnad, cutoff=1):
    """
    Returns the community ID for the reference window that has the largest
    intersection with the matching community at the matching window.
    Communities at the reference window with less than *cutoff* percent of nodes
    are ignored.
    """
    
    trgtComm = -1
    intersectSize = 0
    for commID in dnad.nodesComm[refWindow]["commOrderSize"]:
        # Skip community if it has less than one percent of the nodes.
        commSize = len(dnad.nodesComm[refWindow]["commNodes"][commID])
        if commSize < cutoff:
            continue
        
        tmpSize = len( set(dnad.nodesComm[refWindow]["commNodes"][commID]).intersection( 
            set(dnad.nodesComm[mWindow]["commNodes"][mCommID]) ) )
        
        # Selects the largets intersection
        if intersectSize < tmpSize:
            intersectSize = tmpSize
            trgtComm = commID
    
    return trgtComm, intersectSize

print("Using reference window {0} with highest modularity {1:<1.4}".format(*windModul[0]))

communities = defaultdict(list)
for window in range(dnad.numWinds):
    for commID in dnad.nodesComm[window]["commOrderSize"]:
        
        # Skip community if it has less than one percent of the nodes.
        commSize = len(dnad.nodesComm[window]["commNodes"][commID])
        if commSize < cutoff:
            continue
        
        matchID, interSize = matchComm(commID, window, refWindow, dnad, cutoff)
        
        communities[matchID].append( (commID, interSize, window) )
        
communities = {key:val for (key,val) in communities.items() }
communities.keys()

# Creates a list of communities ID from the dictionary keys
# Orders the keys according to mean intersection size over all windows.
tmpList = []
for key,val in communities.items():
    tmpList.append((key, np.mean([pair[1] for pair in val]), len(val)))
tmpList.sort(key=lambda x:x[1], reverse=True)
tmpList

# Creates a pandas data frame for plotting and analysis
commList = []
genCommID = 0
for key in [x[0] for x in tmpList]:
    val = communities[key]
    for valList in val:
        commList.append( [genCommID, *valList ] )
    genCommID += 1

commDF = pd.DataFrame(data=commList, columns=["genCommID","commID","interSize","Window"])

# %% [code]
# Changes "genCommID" for communities that are matched to the same community in the reference window.
c = commDF.groupby(["genCommID","Window"]).cumcount()
c *= 0.1
commDF[ "genCommID" ] += c

# %% [code]
# Creates a NumPy 2D array to organize data and transform it in a pandas DF.
# Not pretty but its pynthon...
nodeCommNP = np.empty([dnad.numNodes, dnad.numWinds])
nodeCommNP.fill(-1)

#Group by general community ID
grpBy = commDF.groupby("genCommID")
for genCommID, group in grpBy:
    for winIndx,commID in group[["Window","commID"]].values:
        for node in range(dnad.numNodes):
            if dnad.nxGraphs[winIndx].nodes[node]["modularity"] == commID:
                nodeCommNP[node, winIndx] = genCommID

# %% [code]
# Removes nodes that were not classified in a "big-nough" (bigger than 1%) cluster in *any* window.
nodeCommDF = pd.DataFrame(data=nodeCommNP,columns=["Window"+str(i) for i in range(dnad.numWinds)])
nodeCommDF["Node"] = [i for i in range(dnad.numNodes)]
nodeCommDF = nodeCommDF[ nodeCommDF.min(1) >= 0]
# So we don't get "blank"/empty areas in the plot
nodeCommDF["NodePlot"] = [i for i in range(len(np.unique(nodeCommDF["Node"])))]

# %% [code]
# Checks that target nodes are classified in ALL windows
nodeCommDF.loc[ nodeCommDF["Node"].isin(trgtNodes) ]

# %% [code]
# Melts for plotting.
nodeCommDFmelt = nodeCommDF.melt(id_vars=["Node","NodePlot"], value_name="Cluster", var_name="Window")
# Makes it easier to plot
nodeCommDFmelt["Cluster"] = nodeCommDFmelt["Cluster"].astype('category')
# Makes it easier to plot
for i in range(dnad.numWinds):
    nodeCommDFmelt.replace("Window"+str(i),i, inplace=True)

# %% [code]
nodeCommDFmelt.loc[nodeCommDFmelt["Node"].isin(trgtNodes)].groupby("Node")["Cluster"].apply(np.unique)

# %% [code]
trgtClusters = np.unique( nodeCommDFmelt.loc[nodeCommDFmelt["Node"].isin(trgtNodes), "Cluster"].values )
trgtClusters

# %% [code]
# Add readable info to nodes
def getTagStr(i):
    # Store atom names for residues with multiple nodes
    if len(getNodeFromSel( getSelFromNode(i, dnad.nodesAtmSel), dnad.nodesAtmSel, dnad.atomToNode)) > 1:
        atmStr = ":" + dnad.nodesAtmSel.atoms[i].name
    else:
        atmStr = ""
        
    retStr = dnad.nodesAtmSel.atoms[i].resname.capitalize() + \
            ":" + str(dnad.nodesAtmSel.atoms[i].resid) + \
            atmStr + \
            "_" + dnad.nodesAtmSel.atoms[i].segid
            
    return retStr

nodeCommDFmelt['resid']     = np.vectorize(getTagStr)(nodeCommDFmelt["Node"])

# %% [code]
# Write data for Ploting (plots from ggplot in R are much better!)
nodeCommDFmelt.to_csv(os.path.join(workDir, "cluster.csv"),index=False)

# %% [code]
# Get all nodes that make contact with target nodes in any window
contactNodes = np.unique( np.where( dnad.corrMatAll[:,trgtNodes,:] > 0 )[2] )
contactNodesTrgts = list(trgtNodes)
for node in contactNodes:
    if len( set(trgtClusters).intersection( 
            set(np.unique(nodeCommDFmelt.loc[ nodeCommDFmelt["Node"] == node].Cluster)) ) ) :
        contactNodesTrgts.append(node)

# Save data to file
pd.DataFrame(contactNodesTrgts, columns=["contactNodesTrgts"]).to_csv(
    os.path.join(workDir, "contactNodesTrgts.csv"),index=False)

pd.DataFrame(trgtNodes, columns=["trgtNodes"]).to_csv(
    os.path.join(workDir, "trgtNodes.csv"),index=False)

# %% [markdown]
### Prepare pandas data frame with community data

# %% [code]
# Prepares variable names for multi-system comparisons. In this tutorial, we only have one system.
system1 = "OMP"
cDF = nodeCommDFmelt
cDF["system"] = system1
refWindow1 = refWindow

# %% [code]
# Loads VMD-compatible color scales to match community colors in R plots, NGLView, and VMD figures.
comColorScale = getCommunityColors()

# %% [code]
%%R -i workDir,refWindow,comColorScale -o colorValues,clusterIDs -w 800 -h 450

dataPath = file.path(workDir, "cluster.csv")

dt <- fread(dataPath)
clusterIDs = dt[, unique(Cluster)]

colourCount = length(unique(dt$Cluster))

# We only have 50 availabl colors
colourCount <- min(colourCount,50)

rgbCodes <- data.table(comColorScale)

colorValues <- sapply(seq(colourCount), function(x) rgb(rgbCodes[x, .(R,G,B) ],  maxColorValue = 255) )

setorder(dt, Cluster)                      
colorValues = setNames(colorValues, dt[, unique(Cluster)])
colorValues           

print(paste("Creating palette for",colourCount,"clusters"))

# %% [code]
# Builds a color dictionary from the cluster color scale built in R
colorValDict = {}
colorValDictRGB = {}
for key,val in zip(clusterIDs, list(colorValues)):
    colorValDict[key] = val

for key,val in colorValDict.items():
    colorValDictRGB[key] = tuple(int(val.lstrip('#')[i:i+2], 16) for i in (0, 2 ,4))

# %% [markdown]
### View community in the structure

# %% [markdown]
#### Connections to the ligand: 
First, we show all direct network connections between the ligand and surounding nodes from the active site. The next **two** jupyter notebook cell should be executed to show a sliding selection widget that will allow you to compare the network architectures of the acitve site in different simulation windows.

Solid lines indicate connections between nodes in the same community, while dashed lines indicate connections between nodes of different communities.

# %% [code]
w = nv.show_mdanalysis(workUviz.select_atoms("all"))

w._remote_call("setSize", target="Widget", args=["800px", "600px"])
w.parameters = dict(theme='light')

w

# %% [code]
windowSlider = widgets.IntSlider(
    value=refWindow, min=0, max=dnad.numWinds-1, step=1, description='Window:', readout_format='d',
    disabled=False, continuous_update=False, orientation='horizontal', readout=True )

def updateCommByTarget(value):
    w.clear_representations()
    w.add_cartoon("protein")
    w.add_hyperball(ligandSegID)
    showCommunityByTarget(w, nodeCommDF, trgtNodes, windowSlider.value, dnad.nodesAtmSel, dnad, colorValDict)
    
windowSlider.observe(updateCommByTarget,  names='value')

windowSlider

# %% [code]
w.close()

# %% [markdown]
#### Entire communities:

Second, we show all nodes from a chosen community. The following jupyter notebook cell different options to choose which community is shown. The first will select a community ID and show all nodes. The second allows you to select a set of nodes and show them in the structure.

# %% [code]
w = nv.show_mdanalysis(workUviz.select_atoms("all"))

w._remote_call("setSize", target="Widget", args=["800px", "600px"])
w.parameters = dict(theme='light')

opac = 1

w.clear_representations()
w.add_cartoon("protein", color='lightgray', opacity=opac)
w.add_hyperball(ligandSegID)

shapeCounter = [0]
w

# %% [code]
# Dysplays all nodes of a cluster and colors them by cluster ID.

clusterID = 0
showCommunityByID(w, cDF, clusterID, system1, refWindow, 
                  shapeCounter, dnad.nodesAtmSel, colorValDictRGB, system1, refWindow1)

# %% [code]
# Dysplays a given list of nodes and colors them by cluster ID.

clusterID = 0
nodeList = cDF.loc[ (cDF.system == system1) & (cDF.Window == refWindow1) & (cDF.Cluster == clusterID) ].Node.values

showCommunityByNodes(w, cDF, nodeList, system1, refWindow1, shapeCounter, dnad.nodesAtmSel, colorValDictRGB)

# %% [code]
# Color ALL nodes by community

nodeList = cDF.loc[ (cDF.system == system1) & (cDF.Window == refWindow1) ].Node.values
showCommunityByNodes(w, cDF, nodeList, system1, refWindow1, shapeCounter, dnad.nodesAtmSel, colorValDictRGB)

# %% [code]
w.close()

# %% [markdown]
### Make more plots: Clustering of nodes

# %% [code]
plotFilePrefix = "OMP_"

# %% [code]
%%R -i workDir,plotFilePrefix -w 800 -h 450

dataPath = file.path(workDir, "cluster.csv")
plotPath = file.path(workDir, paste0("Plots/",plotFilePrefix,"Clusters_Node_vs_Window.png"))

dt <- fread(dataPath)
dt <- dt[,.(NodePlot,Window,Cluster)]
dt <- dt[, Cluster := as.factor(Cluster) ]

p <- ggplot(dt) + 
    geom_raster(aes(x=NodePlot, y=Window, fill=Cluster)) + 
    scale_fill_manual(values = colorValues) +
    labs(x="Node", y="Window") +
    theme_bw(base_size=20)

ggsave(plotPath, p, device="png")

p

# %% [code]
%%R -i workDir,refWindow,trgtNodes,plotFilePrefix -w 800 -h 450

dataPath = file.path(workDir, "cluster.csv")
plotPath = file.path(workDir, paste0("Plots/",plotFilePrefix,"Clusters_Node_vs_Window_Grouped.png"))

dt <- fread(dataPath)
dt <- dt[, Cluster := as.factor(Cluster) ]

setorder(dt, Cluster, Window)
dt <- dt[, NodePlot := as.factor(NodePlot) ]

# Get the actual indices of the nodes in the re-ordered x-axis (grouped by cluster)
trgtIndices = data.table(trgt = c(which(dt[Window == refWindow]$Node %in% trgtNodes))  )

breaksNodePlot = dt[Window == refWindow,][ Node %in% trgtNodes, NodePlot ]

labelsNode = dt[Window == refWindow,][ Node %in% trgtNodes, Node ]

# Build base plot
p <- ggplot(dt) + 
    geom_raster(aes(x=NodePlot, y=Window, fill=Cluster)) 

# Add intercept lines for target nodes
p <- p + geom_vline(data=trgtIndices, aes(xintercept=trgt), alpha=0.8, linetype = "dashed")
p <- p + geom_hline(aes(yintercept=refWindow), alpha=0.9, linetype = "dashed")

# Finish building plot
p <- p + scale_fill_manual(values = colorValues) +
    scale_x_discrete(limits=dt[Window == refWindow]$NodePlot, breaks=breaksNodePlot, labels=labelsNode) +
    labs(x="Nodes", y="Window") +
    theme_bw(base_size=20)

ggsave(plotPath, p, device="png")

p

# %% [code]
%%R -i workDir,refWindow,trgtNodes,contactNodesTrgts,plotFilePrefix -w 800 -h 450

dataPath = file.path(workDir, "cluster.csv")
plotPath = file.path(workDir, paste0("Plots/",plotFilePrefix,"Clusters_Node_vs_Window_Grouped_ActiveSite.png"))

dt <- fread(dataPath)
dt <- dt[,.(Node,Window,Cluster,resid)]
dt <- dt[, Cluster := as.factor(Cluster) ]
dt <- dt[, Node := as.factor(Node) ]
setorder(dt, Cluster, Window)

dt <- dt[ Node %in% contactNodesTrgts, ]

dt <- dt[, resid := sapply(strsplit(resid, "_"), '[', 1) ]

# Build base plot
p <- ggplot(dt) + 
    geom_raster(aes(x=Node, y=Window, fill=Cluster)) 

# Get the actual indices of the nodes in the re-ordered x-axis (grouped by cluster)
trgtIndices = data.table(trgt = c(which(dt[Window == refWindow]$Node %in% trgtNodes))  )

# Add intercept lines for target nodes
p <- p + geom_vline(data=trgtIndices, aes(xintercept=trgt), alpha=0.9, linetype = "dashed")
p <- p + geom_hline(aes(yintercept=refWindow), alpha=0.9, linetype = "dashed")

# Finish building plot
p <- p + scale_fill_manual(values = colorValues) +
    scale_x_discrete(limits=dt[Window == refWindow]$Node, label=dt[Window == refWindow]$resid) +
    labs(x="Nodes", y="Window") +
    theme_bw(base_size=20) + 
    theme(axis.text.x = element_text(angle = 45, vjust = 1, hjust=1))

ggsave(plotPath, p, device="png")

p

# %% [code]
%%R -i workDir,refWindow,trgtNode,trgtNodes,plotFilePrefix,ligandSegID -w 600 -h 450

trgtNode = as.integer(trgtNode)

dataPath = file.path(workDir, "cluster.csv")
plotPath = file.path(workDir, paste0("Plots/",plotFilePrefix,"Clusters_Node_vs_Window_Grouped_",ligandSegID,".png"))

dt <- fread(dataPath)
dt <- dt[,.(Node,Window,Cluster,resid)]
dt <- dt[ Node %in% trgtNodes, ]
dt <- dt[, Cluster := as.factor(Cluster) ]
dt <- dt[, Node := as.factor(Node) ]
dt <- dt[, Window := as.factor(Window) ]
dt <- dt[, resid := sapply(strsplit(resid, "_"), '[', 1) ]
dt <- dt[, resid := as.factor(resid) ]

setorder(dt, Cluster, Node)

# Build base plot
p <- ggplot(dt) + 
    geom_raster(aes(x=Node, y=Window, fill=Cluster))

# Finish building plot
p <- p + scale_fill_manual(values = colorValues) +
    scale_x_discrete(limits=as.factor(trgtNodes), label=dt[ Node %in% as.factor(trgtNodes), unique(resid)]) +
    scale_y_discrete(limits=dt[Node == trgtNode]$Window, labels=NULL) +
    labs(x="Nodes", y="Window") +
    theme_bw(base_size=20) + 
    theme(axis.text.x = element_text(angle = 45, vjust = 1, hjust=1))

ggsave(plotPath, p, device="png")

p

# %% [markdown]
#  Combine data for Interface and Active Site analysis

Combines interface edges with label information for plots.

Creates structure wirh optimal paths and labels for Active Site analysis, shaping information for interaction between ligand and active site residues.

# %% [code]
window = 0

# Gets all pairs of nodes with non-zero correlations
# nonzeroPairs = [(i,j) for i,j in np.asarray(np.where(dnad.corrMatAll[window, :, :] > 0)).T if i < j]
nonzeroPairs = [(i,j) for i,j in np.asarray(np.where(dnad.contactMat > 0)).T if i < j]

# Combines cartesian distance and correlation
# We use the mean cartesian distance (0:mean, 1:SEM, 2:Min, 3:Max)
distType = 0
# carCorMat = [ [i, j, 
#                getCartDist(i,j,dnad.numNodes,dnad.nodeDists, distType), 
#                np.mean(dnad.corrMatAll[window,i,j]) ] 
#              for i,j in nonzeroPairs ]
carCorMat = [ [i, j, 
               getCartDist(i,j,dnad.numNodes,dnad.nodeDists, 0), 
               getCartDist(i,j,dnad.numNodes,dnad.nodeDists, 1), 
               np.mean(dnad.corrMatAll[:,i,j]),
               stats.sem(dnad.corrMatAll[:,i,j])] 
             for i,j in nonzeroPairs if np.mean(dnad.corrMatAll[:,i,j]) > 0 ]

carCorMat = pd.DataFrame(carCorMat, columns=["i","j","Cart","CartSEM","Corr","CorrSEM"])

def interNode(i,j,dnad):
    # Checks if the pair of nodes exists in the "interNodePairs" 2D array.
    return len( ( dnad.interNodePairs == [i, j] ).all(axis=1).nonzero()[0] )

# Adds interface information (true/false)
carCorMat["EdgeType"] = carCorMat.apply( lambda x: "Interface" if interNode(x["i"], x["j"], dnad) 
                                        else "Internal" , axis=1)


# %% [code]
# Initialized data frame with pairs of source node (all nodes in the system) and target node (defined above).

dataTmp = []

window = 0
for window in range(dnad.numWinds):
    
    df = pd.DataFrame(np.asarray([ (i,trgt) for trgt in trgtNodes  for i in range(dnad.numNodes)]),
                 columns=["node", "targets"])
    
    df["distances"]  = df.apply( lambda row: dnad.distsAll[window, row["node"], row["targets"]], axis=1)
    # Selects the mean distance (0:mean, 1:SEM, 2:Min, 3:Max)
    distType = 0 
    df["cdistances"] = df.apply( lambda row: getCartDist(row["node"], row["targets"], dnad.numNodes, 
                                                         dnad.nodeDists, distType), axis=1)
    # Selects the standard error of the mean distance (0:mean, 1:SEM, 2:Min, 3:Max)
    distType = 1
    df["cdistSEM"] = df.apply( lambda row: getCartDist(row["node"], row["targets"], dnad.numNodes, 
                                                         dnad.nodeDists, distType), axis=1)
    df["path"]       = df.apply( lambda row: list(getPath(row["node"], row["targets"], 
                                                          dnad.nodesAtmSel, dnad.preds, win= window)), axis=1)
    df['path_lens']  = df.apply( lambda row: len(row["path"]), axis=1)
#     df["mincdist"]   = df.groupby("node")[["cdistances"]].transform(lambda x: np.min(x) )

    def getTagStr(i):
        return dnad.nodesAtmSel.atoms[i].resname.capitalize() + ":" + str(dnad.nodesAtmSel.atoms[i].resid) + \
                "_" + dnad.nodesAtmSel.atoms[i].segid

    df['resid']     = np.vectorize(getTagStr)(df["node"])

    ptnIXs = dnad.nodesAtmSel.select_atoms("protein").ix_array
    nclIXs = dnad.nodesAtmSel.select_atoms("nucleic").ix_array
    def getType(nodeIndx):

        if dnad.nodesAtmSel.atoms[nodeIndx].ix in ptnIXs:
            return "Aminoacid"

        elif dnad.nodesAtmSel.atoms[nodeIndx].ix in nclIXs:
            return "Nucleotide"

        else:
            return dnad.nodesAtmSel.atoms[nodeIndx].resname

    df["type"] = np.vectorize(getType)(df["node"])

    df["Interface"] = df["node"].apply( lambda x: x in dnad.contactNodesInter)

    # "Cleans" all -1 distances betweeen nodes not connected by any path.
    df.loc[ df["path_lens"] == 0, "distances" ] = 0
    
    df["window"] = window
    
    dataTmp.append(df.copy())
    del df
    
df = pd.concat(dataTmp)
del dataTmp

# %% [code]
# df.head()
df.tail()

# %% [code]
df[ (np.isin(df["node"],getNodeFromSel("segid " + ligandSegID, dnad.nodesAtmSel, dnad.atomToNode))) & (df["window"] == 0) ]

# %% [markdown]
# Add labels to selected residues

Using references form the literature

# %% [code]
df["Label"] = 0

tags = dict()
#"resname H2U and element N"
tags["OMPn"] = getNodeFromSel("resname OMP and name N1", dnad.nodesAtmSel, dnad.atomToNode)[0]
#"resname H2U and element P"
tags["OMPp"] = getNodeFromSel("resname OMP and name P", dnad.nodesAtmSel, dnad.atomToNode)[0]

for label, node in tags.items():
    if not node in df.node:
        print("Skiping node not found:",node)
        continue
    df.loc[ df.node == node, 'Label'] = label


# %% [code]
df.loc[ (df["resid"] == "Omp:301_OMP") & (df["window"] == 0)  ]

# %% [markdown]
### Adds label and type information for all pairs of connected residues in the system

# %% [code]

carCorMat["iRes"] = carCorMat.apply(lambda row: df[ (df["node"] == row["i"]) & 
                               (df["targets"] == trgtNode[0])]["resid"].iloc[0] , axis=1)

carCorMat["jRes"] = carCorMat.apply(lambda row: df[ (df["node"] == row["j"]) & 
                               (df["targets"] == trgtNode[0])]["resid"].iloc[0] , axis=1)

carCorMat["iType"] = carCorMat.apply(lambda row: df[ (df["node"] == row["i"]) & 
                               (df["targets"] == trgtNode[0])]["type"].iloc[0] , axis=1)

carCorMat["jType"] = carCorMat.apply(lambda row: df[ (df["node"] == row["j"]) & 
                               (df["targets"] == trgtNode[0])]["type"].iloc[0] , axis=1)

# %% [code]
carCorMat.head()

# %% [code]
carCorMat.tail()

# %% [code]
# Create a copy only with interface connections, and check that i < j
carCorMatInterface = carCorMat[ carCorMat["EdgeType"] == "Interface" ]

# %% [markdown]
### Compare Intra and Inter segment correlations

# %% [code]
%%R -i carCorMat,workDir -w 800 -h 500

plotPath = file.path(workDir, "Plots/Interf_Intern_Cart_vs_Corr.png")

cartCutoff = 4.0

dt = data.table(carCorMat)
dt  = dt[Cart < cartCutoff]

p <- ggplot(dt) + 
    geom_point( aes(x=Cart, y=Corr, color=EdgeType), alpha=0.7, size=3 ) +
    geom_smooth( aes(x=Cart, y=Corr, color=EdgeType) ) +
    labs(x="Mean Cartesian Distance (A)", y="Mean Correlation", color="Edge Type") +
    scale_color_brewer(type='qual', palette=6) +
    theme_linedraw(base_size=20) + xlim(c(2.5, cartCutoff))

ggsave(plotPath, p, device="png")

p


# %% [markdown]
### Compare network connectivity, network distance and cartesian distance

# %% [code]
# Select all nodes involved in at least one "Interface" connection.
tmpDF = carCorMat[ carCorMat["EdgeType"] == "Interface" ]

tmpNodeSet = set(tmpDF["i"])
tmpNodeSet.update( set(tmpDF["j"]) )

nodeContacs = []
for node in tmpNodeSet:

    tmp = tmpDF.loc[  (tmpDF["i"] == node) | (tmpDF["j"] == node) ]["Corr"]
    tmp2 = tmpDF.loc[  (tmpDF["i"] == node) | (tmpDF["j"] == node) ]["Cart"]
    
    label = df.loc[ df["node"] == node ]["resid"].unique()[0]
    
    nodeContacs.append( [node, int(tmp.size), 
                         tmp.mean(), sp.stats.sem( tmp ),
                         tmp2.mean(), sp.stats.sem( tmp2 ),
                         label] )

nodeContacs = pd.DataFrame(nodeContacs, 
                           columns=["Node", "NumContacts", "MeanCorr", "SEMCorr", 
                                    "MeanCart", "SEMCart", "label"])
nodeContacs = nodeContacs.fillna(0)

# %% [code]
%%R -i nodeContacs -w 800 -h 600

dt <- data.table(nodeContacs)
sapply(dt, class)

ggplot(dt) + 
#     geom_point(aes(x=NumContacts, y=MeanCorr, color=MeanCart), size=3) +
    geom_point(aes(x=NumContacts, y=MeanCart, color=as.double(MeanCorr)), size=3) +
    geom_label_repel(data=dt[NumContacts >= 10 | MeanCart <= 3 | MeanCorr > 0.8 ], 
                    aes(x=NumContacts, y=MeanCart, label=label)) + 
    scale_y_log10() +
    labs(x="Number of Contacts", 
         y="Mean Cartesian Distance",
         color="Mean Correlation") +
    scale_color_gradient(low="blue",high="red") + 
    theme_linedraw(base_size=20)


# %% [markdown]
## Active stite connections
Here we visualize the residues that make direct connections to the ligand, and display their generalized correlation coefficients.

# %% [code]
%%R -i carCorMatInterface,workDir,ligandSegID -w 800 -h 500

plotPath = file.path(workDir, "Plots/Interface_AAres_vs_Corr_NoLabel.png")

corrCutoff = 0.0

dt <- data.table(carCorMatInterface)

iResList <- dt[iType == "Aminoacid" & jType != "TIP3"][Corr > corrCutoff][,iRes]
dt <- dt[ iRes %in% iResList, ]
dt <- dt[jType != "TIP3", .SD[which.max(Corr)] , by=.(iRes,jRes)]

dt <- dt[, iRes := sapply(strsplit(iRes, "_"), '[', 1) ]

colours <- c("Aminoacid" = "red", 
          "TIP3" = "blue", 
          "Nucleotide" = "darkorange")
colours <- c(colours, setNames("purple",ligandSegID))

shapes <- c("Aminoacid" = 19,
          "TIP3" = 15, 
          "Nucleotide" = 17)
shapes <- c(shapes, setNames(19,ligandSegID))

p <- ggplot(dt) + 
    geom_linerange( aes(x=iRes, y=Corr, ymin=Corr-CorrSEM, ymax=Corr+CorrSEM), size=1 )+
    geom_point( aes(x = iRes, y = Corr, color = jType, size=Cart) )+
    labs(x="Aminoacid Residue", y="Mean Correlation", color="Residue Type", size="Mean Distance") + 
    scale_colour_manual(name = "Residue Type",
                     values = colours) +
    guides(colour = guide_legend(override.aes = list(size=5))) +
    theme_classic(base_size = 20) +
    theme(axis.text.x = element_text(angle = 45, vjust = 1, hjust=1),
          panel.grid.major.x = element_line(color="black", size=0.5, linetype="dotted"))

ggsave(plotPath, p, device="png")

p

# %% [markdown]
### Vizualize 

# %% [code]

w = nv.show_mdanalysis(workUviz.select_atoms("all"))
w._remote_call("setSize", target="Widget", args=["800px", "600px"])
w.parameters = dict(theme='light')
w


# %% [code]
# Paint all connections to the ligand
window = 1

w.clear_representations()
w.add_cartoon("protein")
w.add_hyperball(ligandSegID)

tmpDF = carCorMat[(carCorMat["i"].isin(trgtNodes)) | (carCorMat["j"].isin(trgtNodes))]
tmpDF = tmpDF.loc[carCorMat["EdgeType"] == "Interface"]

for pair in tmpDF[["i","j"]].values:
    viewPath(w, getPath(pair[0], pair[1], dnad.nodesAtmSel, dnad.preds), 
             dnad.distsAll, dnad.maxDirectDist, dnad.nodesAtmSel, window)
    w.add_ball_and_stick(getNGLSelFromNode(pair[0], dnad.nodesAtmSel, atom=True))
    w.add_ball_and_stick(getNGLSelFromNode(pair[1], dnad.nodesAtmSel, atom=True))

# %% [code]
w.close()

# %% [markdown]
# Pretty Figures - Preparing files for VMD

The next few steps will define the files that will be created for loading into VMD, a popular molecular visualization software.

**Important:**

Make sure you have all the files created for VMD. For instance, if you are not interested in allosteric communications and suboptimal paths, just select two arbitrary nodes so that the path files are created correctly. Alternatively, you may want to adapt the tcl scripts provided at the end of the notebook to your particular case. 


# %% [code]
import copy

# This will be used to look for the maximum and  minimum betweenness value in the graph.
# The maximum value will be used ot normalize all betweenness values for better vizualization.
# The minimum value will be used in case a betweenness value could not be assigned for a given edge,
#    also helping visualization.

# Initialize variable with high value.
minimumBetweeness = 100
# Initialize variable with low value.
maximumBetweeness = -1

for pair,btw in dnad.btws[winIndx].items():
    if btw < minimumBetweeness:
            minimumBetweeness = btw
    if btw > maximumBetweeness:
            maximumBetweeness = btw

# Normalize the value.
minimumBetweeness /= maximumBetweeness
            
for winIndx in range(dnad.numWinds):

    normCorMat = copy.deepcopy( dnad.corrMatAll[winIndx,:,:] )
    normCorMat /= normCorMat.max()
    
    ##########################################################################################
    ### Create PDB file with the system in the first step of each window, for VMD vizualization.
    
    pdbVizFile = os.path.join(workDir, 
                            "networkData_Structure_window_{}.pdb".format(winIndx))
    
    # Calculate number of frames per window.
    winLen = int(np.floor(workUviz.trajectory.n_frames/dnad.numWinds))
    
    # Positions the trajectory at the middle of each window.
    workUviz.trajectory[(winIndx+1)*round(winLen/2)]
    
    with mda.Writer(pdbVizFile, multiframe=False, bonds="conect", n_atoms=workUviz.atoms.n_atoms) as PDB:
        PDB.write(workUviz.atoms)
    
    ##########################################################################################
    ### Create network data file with ALL edges and their normalized weights.
    
    fileName = os.path.join(workDir, 
                            "networkData_AllEdges_window_{}.dat".format(winIndx))
    with open(fileName, "w") as outfile:

        for pair in np.asarray( np.where( np.triu(normCorMat[:,:]) ) ).T:

            node1 = pair[0]
            node2 = pair[1]

            # Get VMD indices for the atoms
            pdbIndx1 = dnad.nodesAtmSel.atoms[node1].id -1
            pdbIndx2 = dnad.nodesAtmSel.atoms[node2].id -1

            string = "{} {} {}".format(pdbIndx1, pdbIndx2, normCorMat[ node1, node2])
            
            outfile.write( string + "\n" )
    
    
    ##########################################################################################
    ### Create network data file with ALL NODES, the maximum normalized weight of edges it belongs to,
    ### and the community it belongs to.
    
    fileName = os.path.join(workDir, 
                            "networkData_AllNodes_window_{}.dat".format(winIndx))
    with open(fileName, "w") as outfile:
        
        for node1 in range(dnad.numNodes):
            
            # Get the VMD index for the atom
            pdbIndx1 = dnad.nodesAtmSel.atoms[node1].id -1
            
            # Get the community the node belongs to
            community1 = int(nodeCommNP[node1, winIndx])
            
            # Find the node to which "node1" is connected with highest correlation.
            node2 = np.where( normCorMat[node1,:] == normCorMat[node1,:].max() )[0][0]
            
            # Skip nodes not assigned to any community
            if community1 < 0:
                continue
            
            string = "{} {} {}".format(pdbIndx1, normCorMat[ node1, node2], community1)
            
            outfile.write( string + "\n" )
    
    
    ##########################################################################################
    ### Create network data file with INTRA-COMMUNITY edges and their normalized weights.
    
    fileName = os.path.join(workDir, 
                            "networkData_IntraCommunities_window_{}.dat".format(winIndx))
    with open(fileName, "w") as outfile:
        
        for pair in np.asarray( np.where( np.triu(normCorMat[:,:]) ) ).T:
            
            node1 = pair[0]
            node2 = pair[1]

            # Checks if both nodes belong to the same community. 
            # If they don't, skip this edge. We only write intra-community edges in this file!
            if nodeCommNP[node1, winIndx] != nodeCommNP[node2, winIndx] :
                continue
            
            # If both nodes do not belong to any community (assigned to community -1), also skip the edge.
            if nodeCommNP[node1, winIndx] < 0:
                continue
            
            community1 = int(nodeCommNP[node1, winIndx])
            
            # Get VMD indices for the atoms
            pdbIndx1 = dnad.nodesAtmSel.atoms[node1].id -1
            pdbIndx2 = dnad.nodesAtmSel.atoms[node2].id -1

            string = "{} {} {} {}".format(pdbIndx1, pdbIndx2, normCorMat[ node1, node2], community1)
            
            outfile.write( string + "\n" )
    
    
    ##########################################################################################
    ### Create network data file with INTER-COMMUNITY edges and their normalized weights.
    
    fileName = os.path.join(workDir, 
                                "networkData_InterCommunities_window_{}.dat".format(winIndx))
    with open(fileName, "w") as outfile:
        
        for pair in np.asarray( np.where( np.triu(normCorMat[:,:]) ) ).T:
            
            node1 = pair[0]
            node2 = pair[1]

            # Checks if both nodes belong to the same community. 
            # If they don't, skip this edge. We only write intra-community edges in this file!
            if nodeCommNP[node1, winIndx] == nodeCommNP[node2, winIndx] :
                continue
            
            # If either node does not belong to any community (assigned to community -1), also skip the edge.
            if (nodeCommNP[node1, winIndx] < 0) or (nodeCommNP[node2, winIndx] < 0):
                continue
            
            community1 = int(nodeCommNP[node1, winIndx])
            community2 = int(nodeCommNP[node2, winIndx])
            
            # Get VMD indices for the atoms
            # VMD uses a 0-based index, so we subtract 1 from the PDB index
            pdbIndx1 = dnad.nodesAtmSel.atoms[node1].id -1
            pdbIndx2 = dnad.nodesAtmSel.atoms[node2].id -1

            string = "{} {} {} {} {}".format(pdbIndx1, pdbIndx2, 
                                             normCorMat[ node1, node2], community1, community2)
            
            outfile.write( string + "\n" )
    
    
    ##########################################################################################
    ### Create file with edges listed by betweeness value (highest to lowest).
    
    fileName = os.path.join(workDir, 
                            "networkData_Betweenness_window_{}.dat".format(winIndx))
    with open(fileName, "w") as outfile:
        
        for pair,btw in dnad.btws[winIndx].items():
            
            node1 = pair[0]
            node2 = pair[1]

            # If either node does not belong to any community (assigned to community -1), also skip the edge.
            if (nodeCommNP[node1, winIndx] < 0) or (nodeCommNP[node2, winIndx] < 0):
                continue
            
            community1 = int(nodeCommNP[node1, winIndx])
            community2 = int(nodeCommNP[node2, winIndx])
            
            # Get VMD indices for the atoms
            # VMD uses a 0-based index, so we subtract 1 from the PDB index
            pdbIndx1 = dnad.nodesAtmSel.atoms[node1].id -1
            pdbIndx2 = dnad.nodesAtmSel.atoms[node2].id -1

            string = "{} {} {} {} {} {}".format(pdbIndx1, pdbIndx2, 
                                                normCorMat[ node1, node2], btw/maximumBetweeness, 
                                                community1, community2)
            
            outfile.write( string + "\n" )


# %% [markdown]
### Write Optimal and Sub-Optimal Paths

Using the convenience functions "getNodeFromSel" and "getSelFromNode", one can easily probe the system and determine the relationship between node in the network graph and the atoms and residues they represent in the actual system.

- getSelFromNode
- getNodeFromSel

See examples below:

# %% [code]
srcNode = getNodeFromSel("resname VAL and resid 11 and segid ENZY",dnad.nodesAtmSel, dnad.atomToNode)[0]
print("Source node:", srcNode)

trgNode = getNodeFromSel("resname ALA and resid 69 and segid ENZY",dnad.nodesAtmSel, dnad.atomToNode)[0]
print("Target node:", trgNode)

# %% [code]
getSelFromNode(58,dnad.nodesAtmSel)

# %% [markdown]
**Create a list of important paths:**

Once you have chosen the nodes that define each path of interest, create a list in the cell below with the indices of the source and target nodes.

To make sure that the VMD scripts will run without the need to be adapted, the user must select at least one pair of nodes. 

# %% [code]
# For example, to write the paths between node 0 (Valine 11) and nodes 58 and 60, the following
#   list must be created:  
# nodesForPaths = [ [0,58], [0,60] ]
#

nodesForPaths = [ [0,58], [0,60] ]

# %% [code]

# Determine how many extra sub-optimal paths will be written.
numSuboptimalPaths = 5

pathListFile = open(os.path.join(workDir, "paths.list"), "w")

for srcNode, trgNode in nodesForPaths:
    
    tmpList = getSelFromNode(srcNode,dnad.nodesAtmSel, atom=True).split()
    srcNodeSel = "".join([tmpList[1],tmpList[4],tmpList[10]])

    tmpList = getSelFromNode(trgNode,dnad.nodesAtmSel, atom=True).split()
    trgNodeSel = "".join([tmpList[1],tmpList[4],tmpList[10]])
    
    # Adds the path suffix to the file
    pathListFile.write("_{}_{}\n".format(srcNodeSel, trgNodeSel))
    
    for winIndx in range(dnad.numWinds):

        normCorMat = copy.deepcopy( dnad.corrMatAll[winIndx,:,:] )
        normCorMat /= normCorMat.max()

        ##########################################################################################
        ### Create file with edges listed by betweeness value (highest to lowest).
        
        # File name is created based on selections, not node index, for readability.
        
        
        
        fileName = os.path.join(workDir, 
                                    "networkData_Paths_window_{}_{}_{}.dat".format(winIndx, 
                                                                                srcNodeSel, trgNodeSel))
        with open(fileName, "w") as outfile:

            allPaths = []

            # Reconstructs the optimal path from Floyd-Warshall algorithm
            pathFW = nx.reconstruct_path(srcNode, trgNode, dnad.preds[winIndx])

            allPaths.append(pathFW)

            # Behind the scenes, use Dijkstra algorithm to find sub-optimal paths
            for pathSO in islice(nx.shortest_simple_paths(dnad.nxGraphs[0], 
                                                srcNode, trgNode, weight="dist"), 1, numSuboptimalPaths + 1):
                allPaths.append(pathSO)

            # Create a counter of number of paths that go though each edge, among all (sub-)optimal path(s).
            pathCounter = defaultdict(int)
            for pathIndx, pathIter in enumerate(allPaths):
                # Iterate over edges in the path
                for i in range(len(pathIter)-1):

                    node1 = pathIter[i]
                    node2 = pathIter[i+1]

                    pathCounter[(node1, node2)] += 1

            # Normalize the count
            maxCount = np.max(list(pathCounter.values()))
            for pair, count in pathCounter.items():
                pathCounter[pair] = count/maxCount

            for pathIndx, pathIter in enumerate(allPaths):
                # Iterate over edges in the path
                for i in range(len(pathIter)-1):

                    node1 = pathIter[i]
                    node2 = pathIter[i+1]

                    # Get the community each node belongs to
                    community1 = int(nodeCommNP[node1, winIndx])
                    community2 = int(nodeCommNP[node2, winIndx])

                    # If either node does not belong to any community (assigned to community -1), 
                    #     also skip the edge.
                    if (community1 < 0) or (community2 < 0):
                        continue

                    # Get the betweeness value
                    try:
                        btw = dnad.btws[winIndx][( node1, node2)]
                    except:
                        # If one could not be calculated (very few paths going though this edge)
                        # set an arbitrarily low value.
                        btw = minimumBetweeness

                    # Get VMD indices for the atoms
                    # VMD uses a 0-based index, so we subtract 1 from the PDB index
                    pdbIndx1 = dnad.nodesAtmSel.atoms[node1].id -1
                    pdbIndx2 = dnad.nodesAtmSel.atoms[node2].id -1

                    string = "{} {} {} {} {} {}".format(pdbIndx1, pdbIndx2, 
                                                     normCorMat[ node1, node2], 
                                                     btw/maximumBetweeness, pathCounter[(node1, node2)], 
                                                     pathIndx)

                    outfile.write( string + "\n" )

pathListFile.close()

# %% [code]
# Write TCL files to load all data we just wrote down and visualize in VMD.
prepTclViz("networkData", str(dnad.numWinds), ligandSegID, workDir)

# %% [markdown]
# Rendering high-quality images with VMD - Step 3

In the third step, the user can load files produced by the Jupyter notebooks into VMD. An easy to load script will handle all the work, creating a simple graphical  user  interface  (GUI)  that can be used to  easily  render publication-quality  images.   These  renderings  can  represent many aspects of the biomolecular system.

There are *two options* to load the system into VMD and open the GUI:

**1. From inside VMD.**

    After opening VMD, in the VMD Main window, go to: Extensions > Tk Console. 
    
    In the Tk Console, type the following command to navigate to the folder where the Analysis Results were saved (same as in the beginning of this tutorial Step 2): 
    
`cd << path to folder >>`
    
    In the results folder, type the following command to load the results and the GUI: 
    
`source network_view_2.tcl`

**2. When loading VMD.**

    If you choose to load VMD from a terminal (command line) window, navigate to the folder where the Analysis Results were saved (same as in the beginning of this tutorial Step 2): 
    
`cd << path to folder >>`
    
    Load VMD with the following command: 
    
`vmd -e network_view_2.tcl`


# %% [markdown]
### How to use the Network View 2 GUI

The Network View 2.0 GUI was created in a way that allows for easy interaction without deep background knowledge on VMD. If you are an expert VMD user, you can still change anything in the representation, but the GUI might erase your changes when loading some of the specialized features.

The GUI allows user to visualize and render all the properties of the network presented in the Step 2. For instance:

**1.** To visualize the communities just click on "All Communities". You can navigate diferent windows by clicking in the window "step" at the top-left corner. 

**2.** To load the betweenness just click on "Betweenness". You can, at the same time, visualize the communities of the protein nodes, by clicking on "Show/Hide Colors" in the "Color Protein by Communities" tab (only works for proteins).

**3.** The "Representations" tab allows the user to show or hide parts of the structure.

**4.** If you want to render the network in a higher or lower resolution, the "Network Drawing Resultion" tab provides options. Note that after selecting a new resolution, you must load the network representation again.

**5.** Three options are available for quickly start rendering. The first two are GPU-only, and therefore depend on your computer having compatible GPU hardware. The third option uses CPU and should run in most computers. If you want to use a different rendering option, in the VMD Main window, go to: File > Render. Then, select the desired rendering option. **Attention** the quick menu for rendering will always save files with the same name. If you want to render multiple figures, and keep all of them, rename the just-rendered images to avoid overwriting the files. 

# %% [markdown]
# ---- The End ----
```
