## Supplementary figures and images for "Generalized correlation-based dynamical network analysis: a new high-performance approach for identifying allosteric communications in molecular dynamics trajectories"

### nodeAtomGroup.png

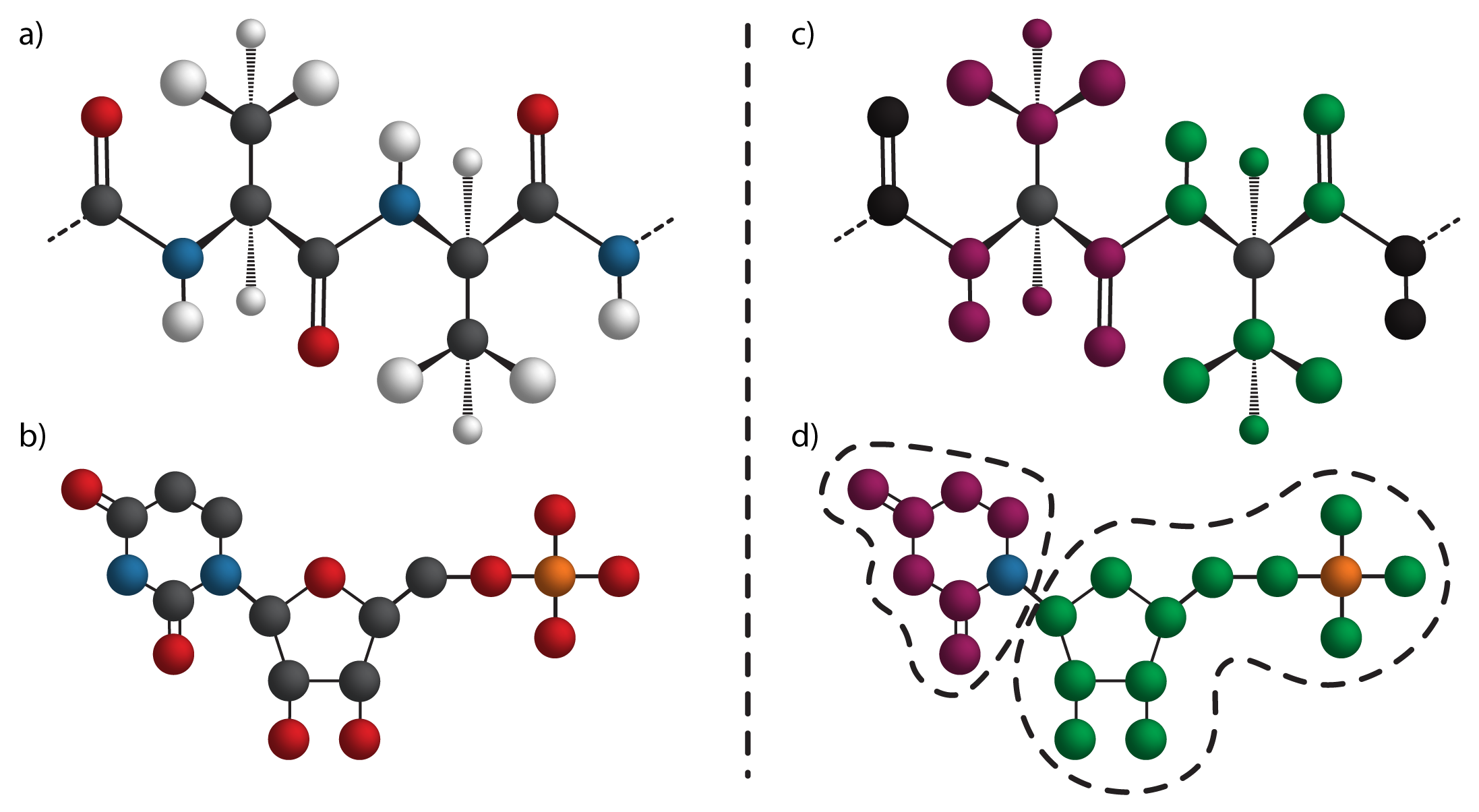

### nodeContactDistances.png

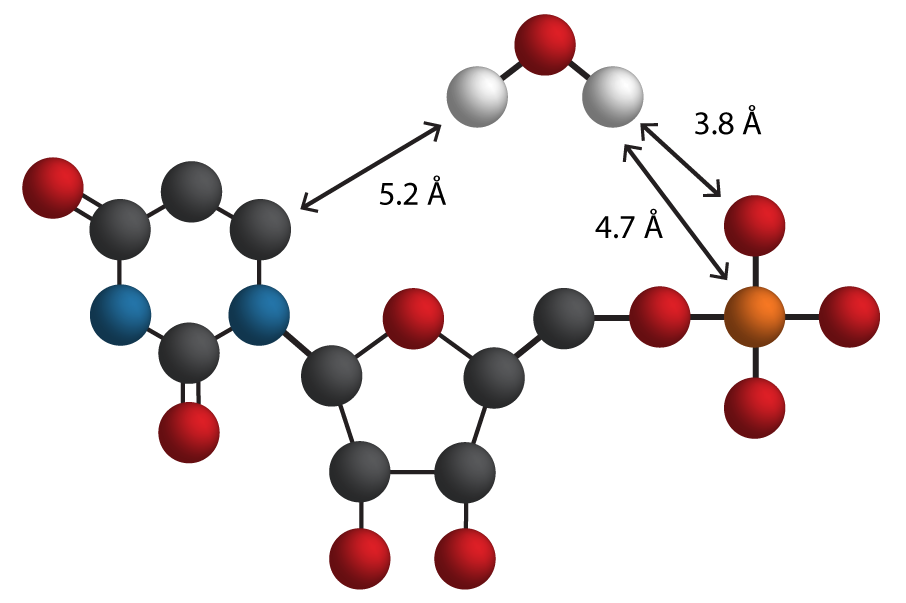
